# Sweet and fatty symbionts: photosynthetic productivity and carbon storage boosted in microalgae within a host

**DOI:** 10.1101/2023.12.22.572971

**Authors:** A. Catacora-Grundy, F. Chevalier, D. Yee, C. LeKieffre, N. L. Schieber, Y. Schwab, B. Gallet, P.H. Jouneau, G. Curien, J. Decelle

## Abstract

Symbiosis between a host and intracellular eukaryotic microalgae is a widespread life strategy in aquatic ecosystems. This partnership is considered to be mainly energized by the photosynthetically-derived carbon energy of microalgal symbionts. A major question is whether microalgae increase their photosynthetic production and decrease carbon storage in order to maximize carbon translocation to their host. By combining three-dimensional subcellular imaging and physiological analyses, we show that the photosynthetic machinery (chloroplast and CO_2_-fixing pyrenoid) of the symbiotic microalga *Micractinium conductrix* significantly expands inside their host (the ciliate *Paramecium bursaria*) compared to the free-living state. This is accompanied by a 13-fold higher quantity of Rubisco enzymes and 16-fold higher carbon fixation rate. Time-resolved subcellular imaging revealed that photosynthetically-derived carbon is first allocated to starch during the day, with five times higher production in symbiosis despite low growth. Nearly half of the carbon stored in starch is consumed overnight and some accumulates in lipid droplets, which are 20-fold more voluminous in symbiotic microalgae. We also show that carbon is transferred to the host and hypothesize that much of this is respired by the high density of surrounding host mitochondria. We provide evidence that the boosted photosynthetic production of symbiotic microalgae could be explained by the energetic demands of the host. Overall, this study provides an unprecedented view of the subcellular remodeling and dynamics of carbon metabolism of microalgae inside a host, highlighting the potentially key role of the source-sink relationship in aquatic photosymbiosis.

**Significance statement:** Symbiotic interactions between a heterotrophic host and intracellular microalgae are widespread in aquatic ecosystems and are considered to be energized by the photosynthetically-derived carbon energy. However, little is known on the impact of symbiosis on the algal bioenergetics (e.g. carbon production and storage). This study reveals the morphological and physiological changes of a microalga inside a host at the subcellular scale over the day. We show that the photosynthetic machinery expands and carbon fixation and storage are boosted in symbiotic microalgae beyond their growth needs. This high photosynthetic production is very likely enhanced by the host energetic demands. Our findings advance our basic understanding of photosymbiosis and open new perspectives on the mechanisms and drivers of metabolic exchange between partners.

## Introduction

Symbiotic associations encompass a broad spectrum of interactions, many of which rely on metabolic exchanges between partners. Photosymbiosis (the association between a heterotrophic host and photosynthetic symbionts) is ubiquitous in aquatic ecosystems. While the most emblematic example of photosymbiosis is the association between corals and microalgae (e.g. Symbiodiniaceae) (Hughes et al. 2017; Sukumaran and T. R. 2023), photosymbiotic interactions with marine and freshwater protists, such as radiolarians, ciliates, dinoflagellates, as hosts are also widespread (Decelle, Colin, and Foster 2015; Stoecker et al. 2009). Although it remains challenging to quantify the benefits for the host and the microalgal symbionts, it is widely considered that photosymbiosis is mutually beneficial, i.e. the host acquires new metabolic capabilities (production of photosynthesis-derived carbohydrates, nitrogen recycling) and symbiotic microalgae benefit from a nutrient-rich environment and protection against predators and viruses (Decelle et al. 2015; Dziallas et al. 2012; Johnson 2011; Yellowlees, Rees, and Leggat 2008). Nevertheless, our mechanistic understanding of this metabolic crosstalk between host and symbiont, particularly the impact of symbiosis on the bioenergetics of microalgae, remains in its infancy. Physiological and morphological changes in symbiotic microalgae have previously been described in unicellular marine plankton photosymbiosis, including expansion of chloroplast volume and higher carbon fixation capability (Decelle et al. 2019, 2021a; Uwizeye et al. 2021). This algal transformation strongly suggests an enhanced primary production of the algae within their hosts, with possible impact of photosymbiosis in global carbon cycles. However, given the wide diversity of taxonomic partners and habitats, it is not known whether this is a common phenomenon in photosymbioses from marine and freshwater ecosystems.

The remodeling of the photosynthetic apparatus in oceanic photosymbiosis raises the question of how microalgae manage their photosynthetically-derived carbon energy within hosts, and more particularly what is the fate of this carbon? In microalgae, carbohydrates (e.g. sugars) produced by photosynthesis are typically used for respiration and growth, or partitioned into storage compounds such as starch (or other glucose polymers) and triacylglycerols (TAG) in lipid droplets (Busi et al. 2014; León-Saiki et al. 2017). Synthesis and degradation of starch and lipids are dynamic (Jouhet et al. 2022; Kong et al. 2018) and depend on cell growth, the time of the day and environmental conditions (León-Saiki et al. 2017; Ran et al. 2019). We speculate that starch and lipid storage in symbiotic microalgae is limited compared to the free-living condition since it has been shown that most (90%) of the organic carbon produced by microalgae is transferred to coral hosts, mainly as glucose and lipids (Davy et al. 2012; Falkowski et al. 1984). Starch and lipids have been observed in microalgae living within corals, Foraminifera and Radiolaria (Decelle et al. 2021b; Gibbin et al. 2020; Krueger et al. 2018), but their diel dynamics inside and outside the host has never been addressed. This knowledge gap prevents us from understanding the impact of the host on the bioenergetics of their symbiotic microalgae, and therefore, the mechanisms and drivers of carbon exchange.

Studying the metabolisms of symbiotic microorganisms is challenging because it requires disentangling the metabolism of both partners with sufficient spatial and temporal resolution. Transcriptomics can provide insights into potentially active metabolic pathways, but do not provide quantitative information (for example about the quantity of carbon that is stored). In this study, we therefore conducted time-resolved 3D ultrastructural imaging to reveal the subcellular architecture of microalgae outside and inside a host over time, quantifying volumes of organelles and compartments that produce and store carbon energy. Combined with physiological measurements, this imaging approach is essential to obtain a full understanding of the metabolic status of a symbiont within a host. We used a single-celled photosymbiotic system, *Paramecium bursaria* (host ciliate) and *Chlorella spp.* (green microalgae, Chlorophyta), that is widely distributed in freshwater ecosystems (Pröschold et al. 2011; Reisser 1980). The ciliate has the capacity to establish a symbiosis with 100-800 algal cells, individually localized in a symbiosome vacuole. These symbiotic microalgae can belong to the genera *Chlorella* and *Micractinium*, which are also able to be cultured in the free-living state (Fujishima 2009). We demonstrate that symbiotic microalgae undergo a major expansion of their chloroplast, including the CO_2_-fixing pyrenoid, which lead to higher carbon fixation compared to the free-living stage. By tracking the fate of this photosynthetically-produced carbon in symbiosis, we show that there is a higher starch production during the day, a higher consumption overnight, carbon accumulation in lipid droplets, and carbon transfer to the host. Our study provides experimental evidence that this high primary productivity could be linked to the energetic demands of the host. Therefore, this study provides new insights into the carbon metabolism of symbiotic microalgae and on potential host processes that engineer carbon energy in photosymbiosis.

## Results and Discussion

### Expanded photosynthetic machinery and higher carbon fixation uptake in symbiotic microalgae

Using the volume electron microscopy technique FIB-SEM (Focused Ion Beam Scanning Electron Microscopy), we compared the subcellular architecture of two free-living microalgae (*Chlorella vulgaris* and *Micractinium conductrix*, both known to be symbionts of the ciliate *Paramecium bursaria*) with that of symbiotic cells (identified here as *M. conductrix*) (Fig. 1). In total, 80 algal cells have been analyzed, representing more than 40,000 electron microscopy images. We focused on 3D reconstruction and quantitative volumetrics of the main algal organelles (i.e. chloroplasts, mitochondria and the nucleus). The two free-living microalgae which are known to engage in symbiosis with *P. bursaria* exhibited similar cellular architecture, with organelles having comparable volumes (Fig. 1B). Having two free-living representatives provides additional evidence on the effect of symbiosis regardless the symbiont taxonomy and culture conditions. By contrast, symbiotic microalgae exhibited significant morphological differences (Fig.1). Cell volume was 5.6-fold higher than the free-living stage (43.55 ± 12.97 µm^3^ *vs* 7.79 ± 1.94 µm^3^). The most important differences involved the energy-producing organelles, with the volume of the chloroplast and mitochondrion increasing 6.4 and 7.3-fold in symbiotic microalgae, respectively (22.92 ± 7.97 µm^3^ *vs* 3.59 ± 1.02 µm^3^ and 1.55 ± 0.52 µm^3^ *vs* 0.21 ± 0.05 µm^3^, respectively) (Fig. 1B). Volume occupancy of the mitochondrion and chloroplast in the cell (organelle/cell volume ratio) tended to be higher in symbiosis (3.52% ± 0.35% in symbiosis *vs* 2.77 ± 0.34 % in free-living and 51.68% ± 4.27% in symbiosis *vs* 45.78% ± 2.18% in free-living, respectively). In contrast, cell volume occupancy of the nucleus was 2-fold lower in symbiosis (5.41 ± 1.14% *vs* 11.18 ± 1.50%) (Fig. 1C). Symbiotic cells exhibited a larger variability of organelle volumes compared to the free-living condition, suggesting different physiological states inside the host. The morphological changes that we observed in this symbiotic microalga share similarities with those reported for some marine planktonic photosymbioses (Decelle et al. 2021a; Uwizeye et al. 2021), suggesting that common cellular processes are involved when photosynthetic production is enhanced within a host.

**Figure 1.**
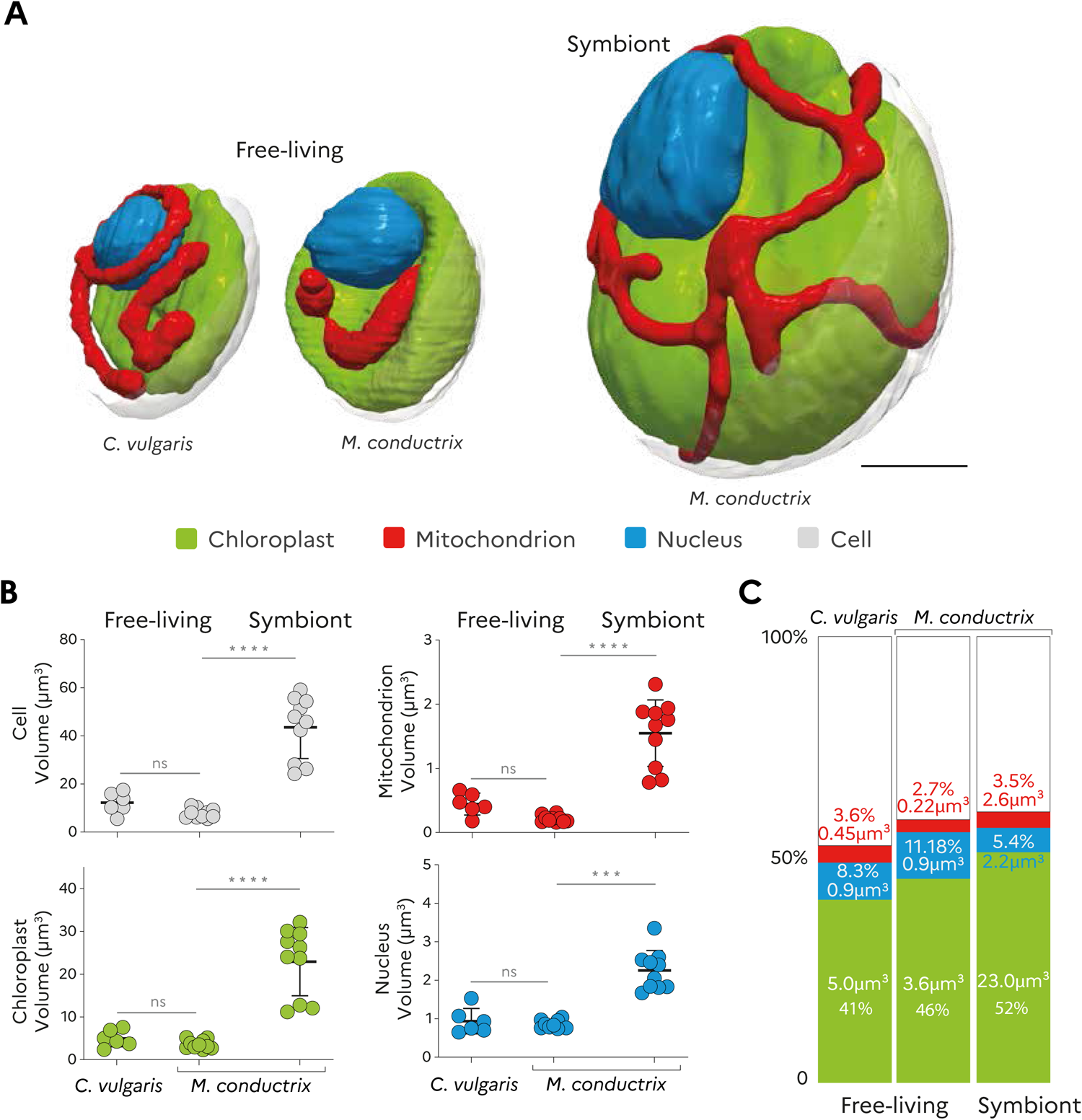
Subcellular architecture of free-living and symbiotic microalgae unveiled by FIB-SEM imaging. **A)** 3D reconstruction of the free-living microalgal cells (*Chlorella vulgaris*; and *Micractinium conductrix*) and the symbiotic microalgal cells (identified as *M. conductrix*) in the host ciliate *Paramecium bursaria* (CCAP1660/18) with chloroplast (green), mitochondrion (red) and the nucleus (blue). Scale bar: 1µm. **B)** FIB-SEM-derived volumes of the cell, chloroplast, mitochondrion and nucleus from the free-living microalgae (*C. vulgaris*; n=6 and *M. conductrix* n=10) and symbiotic microalgae (*M. conductrix*; n=10). Scatter plots present the mean volume of organelles (µm^3^) ± SD. Non-parametric ANOVA unpaired test: ****p < 0.0001 ***p < 0.001; ns, no significant difference. **C)** Relative volume occupancy of the chloroplast, mitochondrion and nucleus in the cell as % (organelle volume/cell volume ratio) in free-living and symbiotic microalgal cells. Volumes (μm^3^) of organelles are given within respective bar segments and summarized in Table S1 to S4. (Grey bar segments represent the remaining volume of the cell).

To investigate the photosynthetic capacities of symbiotic microalgae, we analyzed the structural organization and quantified the volume of the pyrenoid, the CO_2_-fixing liquid-like compartment inside the chloroplast (He, Crans, and Jonikas 2023; Meyer, Whittaker, and Griffiths 2017). 3D reconstruction showed that the pyrenoid matrix of free-living and symbiotic microalgae was traversed by membrane tubules extending from the thylakoids, as previously described in other green microalgae (*Chlorella* and *Chlamydomonas*) (Engel et al. 2015; Fujishima 2009) (Fig. 2A). Compared to free-living cells, the pyrenoid was 8.8-fold larger in symbiosis (1.14 ± 0.27 µm^3^ *vs* 0.13 ± 0.04 µm^3^). It also tended to occupy even more space within the enlarged chloroplast (5.35 ± 1.56% in symbiosis *vs* 3.63 ± 0.70% in free-living) (Fig. 2B). The increased volume of the pyrenoid could contribute to higher carbon fixation by the cell. In order to establish the connection between morphology and function of this key compartment, we quantified carbon fixation (time-resolved incubation with ^13^C-bicarbonate stable isotope) and the content of the CO_2_-fixing enzyme Rubisco (Ribulose-1,5-bisphosphate carboxylase/oxygenase) in free-living and symbiotic microalgae (Fig. 2C-E). Carbon fixation per cell in symbiotic microalgae was 16-fold higher than in free-living cells (0.0192 ± 0.0125 pg of ^13^C.cell^-1^ and 0.0012 ± 0.0002 pg of ^13^C.cell^-1^, respectively) after 1h of incubation with ^13^C-bicarbonate. When normalized per carbon, carbon uptake was 2-fold higher in symbiosis (0.0022 ± 0.00022 pg. ^13^C.cell^-1^ *vs* 0.0011 ± 0.00030 pg. ^13^C.cell^-1^) (Fig. 2C-D – Table S5). Using quantitative western blot, we found that symbiotic cells possess ∼13 times more Rubisco compared to free-living cells (5.77.10^-10^ ± 1.32.10^-10^ pmol.cell^-1^ in free-living *vs* 7.75.10^-09^ ± 4.16.10^-09^ pmol.cell^-1^ in symbiosis) (Fig. 2E-Fig. S1, Table S6), corroborating the increase in pyrenoid volume. This demonstrates that pyrenoid volume assessed by 3D electron microscopy is correlated to Rubisco content, a relationship that has not previously been explored. Higher carbon fixation could also be explained by higher CO_2_ availability surrounding symbiotic microalgae, partly due to the acidification of the symbiosome (Kodama and Fujishima 2005). Overall, the increase in pyrenoid volume and Rubisco content, as well as higher carbon fixation, clearly illustrate significant remodeling of the photosynthetic machinery and enhancement of photosynthetic production by microalgae in symbiosis.

**Figure 2.**
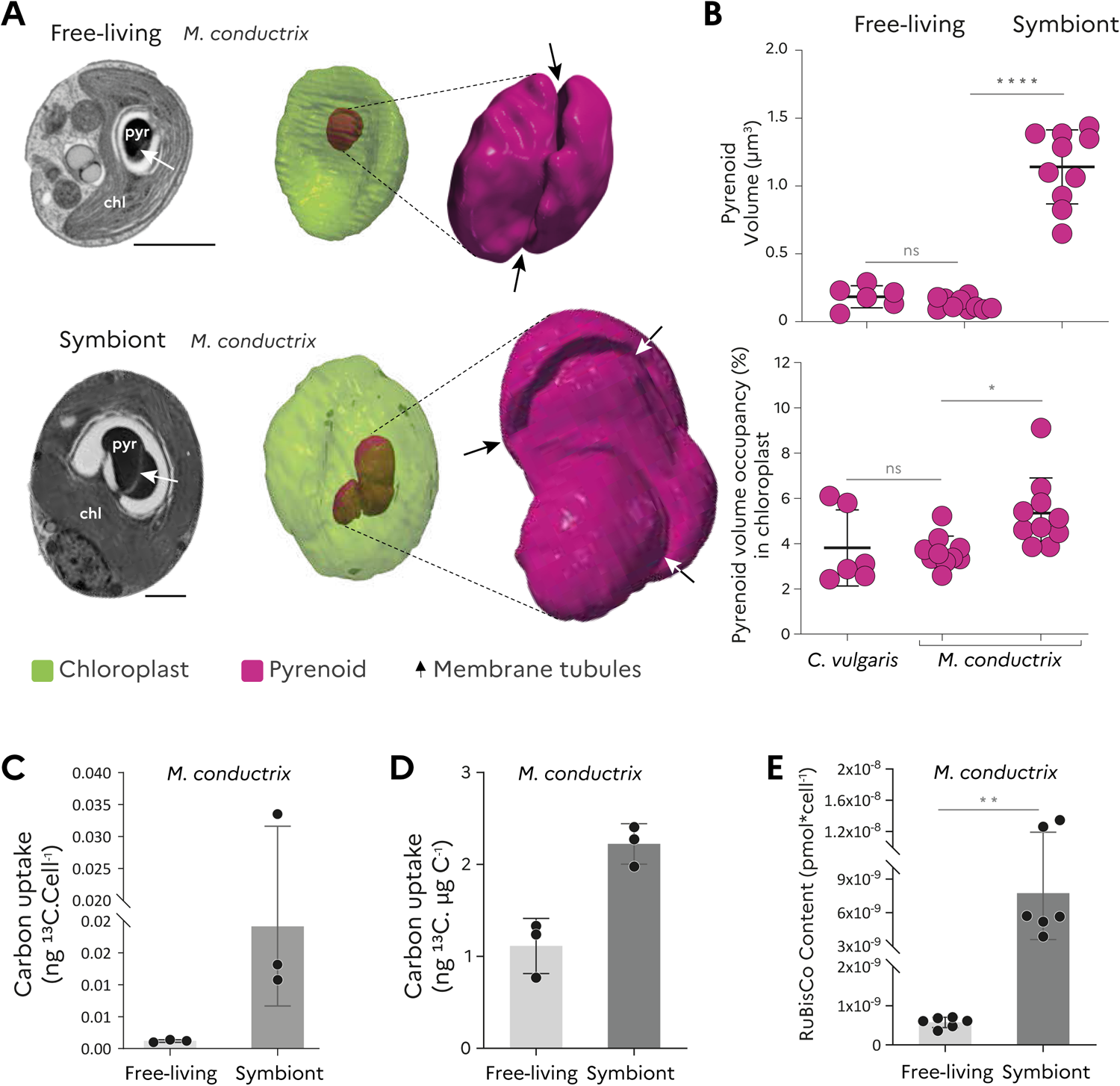
Expansion of the carbon fixation machinery in symbiotic microalgae. **A)** Representative electron micrographs and FIB-SEM-based 3D reconstruction of the chloroplast (chl, green) and its immersed pyrenoid (pyr, purple) in free-living and symbiotic microalgae (*M. conductrix*). Arrows indicate the membrane tubules crossing the pyrenoid matrix. Scale bar: 1 µm. **B)** Scatter plots represent the volume of the pyrenoid in µm^3^ (mean ± SD) and pyrenoid volume occupancy (relative volume of the pyrenoid in the chloroplast) as % (mean ± SD) in free-living *C. vulgaris* (n=6) and *M. conductrix* (n=10) and the symbiotic microalgae (*M. conductrix:* n=10) in the host ciliate *Paramecium bursaria*. Non-parametric ANOVA unpaired test: ****p < 0.0001; *p < 0.01: ns, no significant difference. **C-D)** Inorganic carbon fixation rate after 1h of incubation with ^13^C-labelled bicarbonate in free-living and symbiotic microalgae presented as carbon uptake per algal cell (C) and carbon uptake per carbon (D) in triplicates. **E)** Rubisco content (pmol) per algal cell in free-living and symbiotic state based on Rubisco immunoblot. t-test: **p ≤ 0.05. See also Supplementary Table S6 and Fig S1.

### Higher starch production in symbiotic microalgae and overnight consumption

One of the major questions in photosymbiosis is whether symbiotic microalgae store carbon energy when present inside their host (and if yes, in which quantity) or if this carbon is transferred to the host without storage. In order to investigate carbon allocation in symbiosis, we tracked the fate of the photosynthetically-fixed carbon at the subcellular scale using a correlative TEM-NanoSIMS (Transmission electron Microscopy-Nanoscale Secondary Ion Mass Spectrometry) approach. After 1h of incubation with ^13^C-labelled bicarbonate, ^13^C enrichment was mostly found in starch of the symbiotic microalgae (Fig. 3A), as is the case in symbiotic dinoflagellates from Foraminifera and corals (Kopp et al. 2015; LeKieffre et al. 2018). After 24h, ^13^C enrichment was found not only in starch but also in the algal cytoplasm and algal lipid droplets (Fig. 3B). These results demonstrate that symbiotic microalgae store photosynthetically-fixed carbon during the day in the form of starch in their chloroplast, allocate part of this carbon in newly synthetized biomass for growth, and also store it in lipid droplets, possibly via an overnight reallocation from starch to lipids.

**Figure 3.**
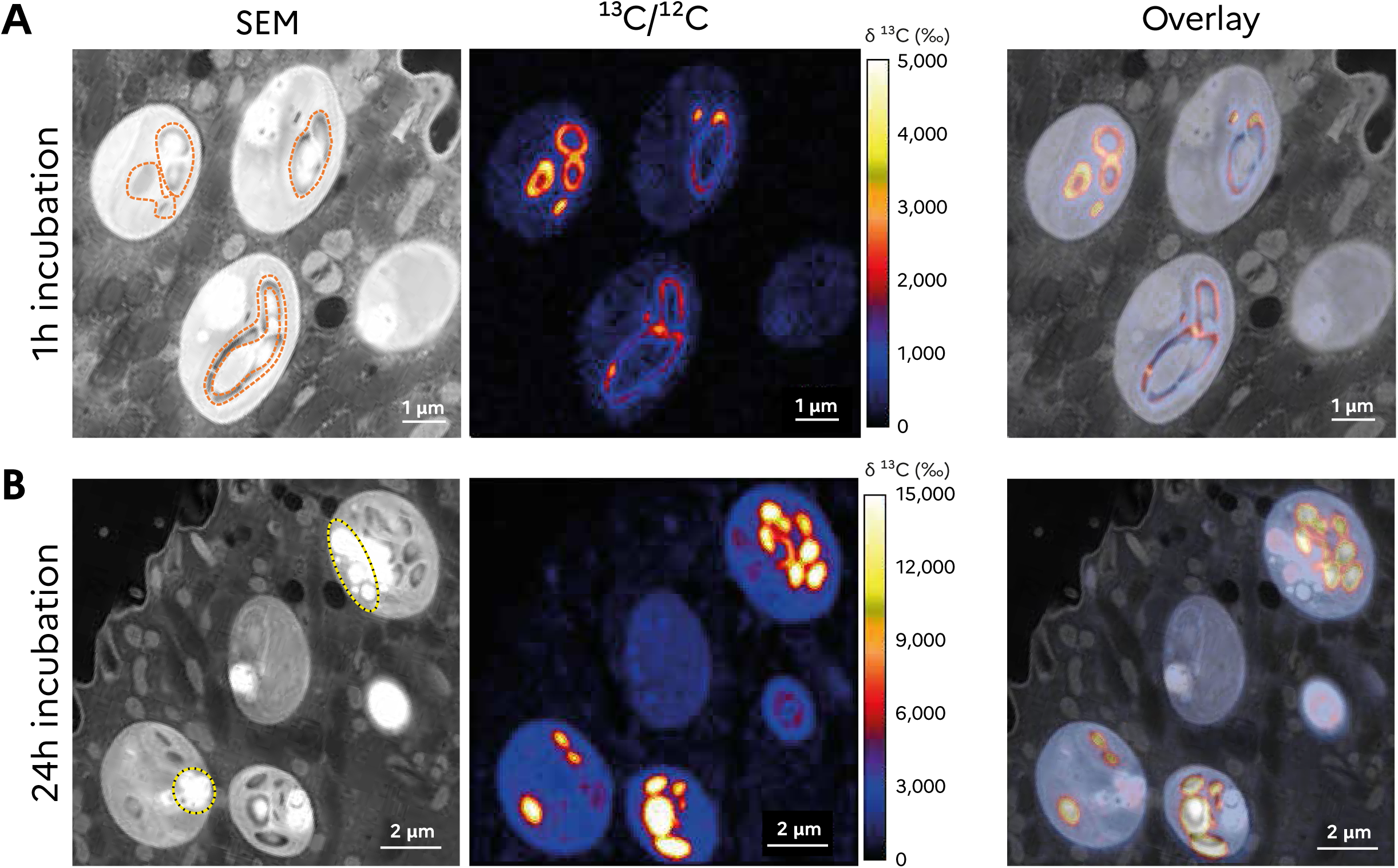
Subcellular tracking of fixed carbon in symbiotic microalgae using mass spectrometry imaging. **A-B)** SEM (Scanning Electron microscopy) and NanoSIMS (Nanoscale Secondary Ion Mass Spectrometry) images and the overlay (right) showing ^13^C enrichment (‰, provided by the ^13^C/^12^C ion map) in symbiotic microalgae within the host *P. bursaria* after 1h (A) and 24h (B) of incubation with ^13^C-labelled bicarbonate. At 1h, ^13^C enrichment was mainly found in starch grains and plates of the symbiotic microalgae while at 24h, ^13^C enrichment was also found in algal lipid droplets. Starch and lipid droplets are highlighted in SEM images by dashed circles in orange and yellow, respectively.

We then investigated whether microalgae store more or less carbon (i.e. starch and triacylglycerols - TAG - in lipid droplets) in symbiosis compared to the free-living stage, and if this storage follows the same temporal dynamics. We combined a bulk quantification of total starch and neutral lipids with FIB-SEM-based volumetrics in free-living and symbiotic microalgae over the day (morning 10 am after 2h of light *versus* afternoon 4pm after 8h of light). Starch quantification in microalgae revealed that synthesis occurs during the day outside and inside a host following the same diel dynamics (Fig. 4A). However, starch production was significantly higher in symbiosis, leading to a 4.4-fold higher starch content per cell at the end of the day (1.40 ± 0.08 pg.cell^-1^ in symbiosis in respect to 0.31 ± 0.12 pg.cell^-1^). It is known that starch production is correlated to cell growth and energetic demands in green microalgae (Ball et al. 1990; Busi et al. 2014). In free-living microalgae, we showed that starch production mainly takes place during exponential growth phase, while at stationary phase, starch content is similar between morning and afternoon (no production at low growth rate, Fig. S2). Despite the 2-times lower growth rate of microalgae in symbiosis (Table S8), as previously reported (Takahashi 2016), microalgae produced 5.3-times more starch during the day (0.91 pg.cell^-1^ in symbiosis *vs* 0.17 pg.cell^-1^ in free-living algae) compared to free-living cells. This may indicate that symbiotic microalgae produce more starch than needed for their growth.

**Figure 4.**
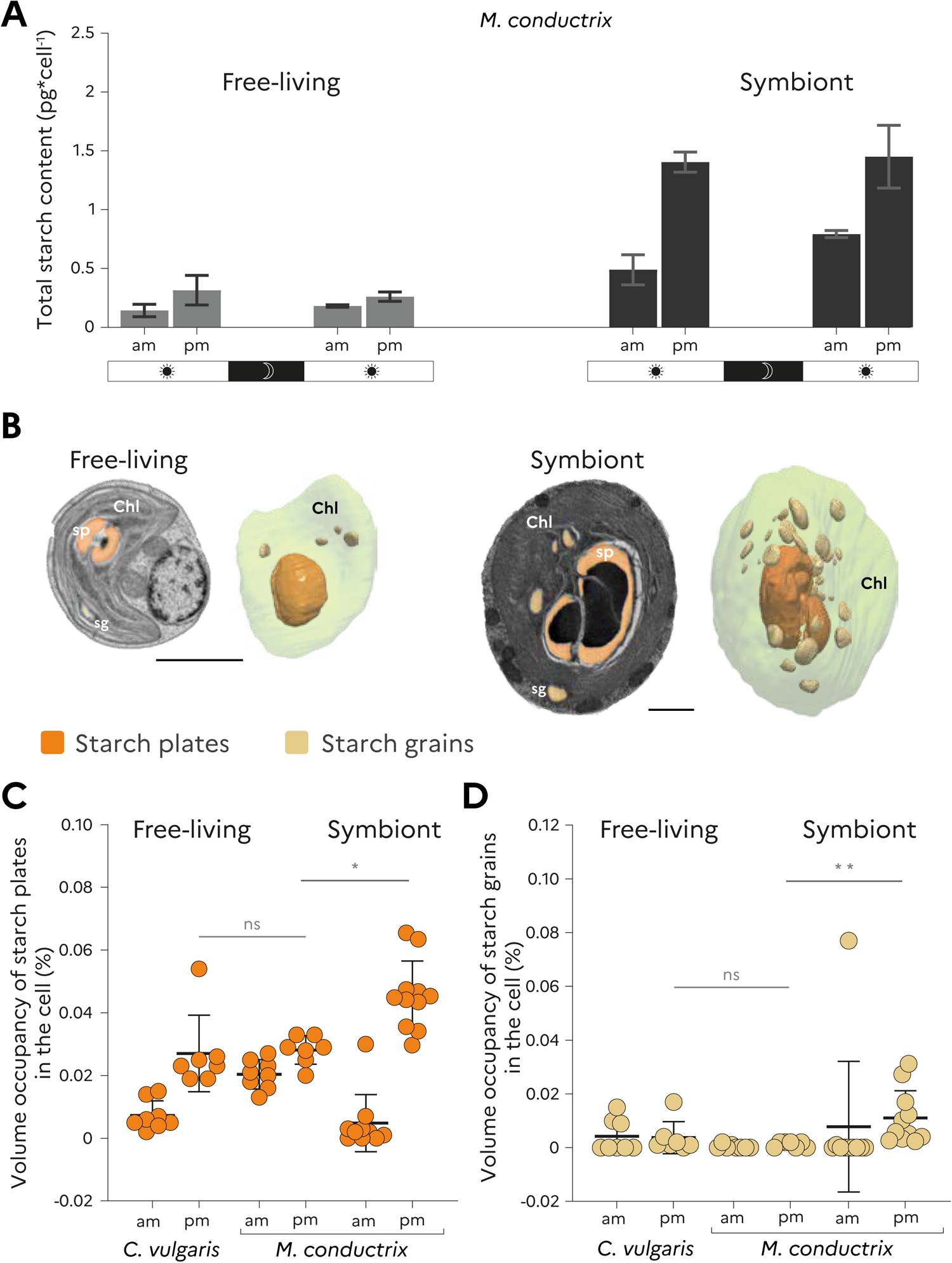
Quantity and diel dynamics of starch synthesis and storage in free-living *versus* symbiotic microalgae. **A)** Starch content per cell in free-living (*C. vulgaris* and *M. conductrix*) and symbiotic microalgae (*M. conductrix*) unveiled by enzymatic assay (n = 3) in the morning (10am) and afternoon (4pm) across two consecutive days. **B)** Electron micrographs and FIB-SEM-based 3D reconstruction of starch plates (sp, dark orange) and starch grains (sg, light orange) in the chloroplast (chl) of free-living and symbiotic microalgae. Scale bar: 1 µm. **C-D)** Scatter plots represent the relative volume of starch plates (C) and starch grains (D) in free-living and symbiotic microalgae (%) provided by FIB-SEM based volumetrics. Non-parametric ANOVA unpaired test: **p<0.01: *p ≤ 0.05: ns, no significant difference. Size scales: 1 µm. See also Supplementary Tables S9 and S10.

FIB-SEM-based 3D reconstruction allowed us to track and quantify two types of starch compartments that are produced depending on growth conditions: starch plates surrounding the pyrenoid and starch grains localized in the stroma of the chloroplast (Fig. 4B) (both labelled with ^13^C with NanoSIMS, Fig. 3). In green microalgae (*Chlamydomonas* sp.) actively dividing cells accumulate more starch in plates, while transitory starch grains can massively increase under stress conditions (Findinier et al. 2019; He et al. 2023) Starch plates are also essential for carbon fixation and the carbon concentrating mechanism (CCM), potentially acting as an oxygen barrier for Rubisco (Toyokawa, Yamano, and Fukuzawa 2020). Compared to free-living cells, FIB-SEM-based morphometrics confirmed the higher amount of total starch per cell volume in symbiotic microalgae at the end of the day (by 2-fold: 0.057 % ± 0.02 *vs* 0.030 % *±* 0.00 per cell volume) (Fig. S3). Specifically, FIB-SEM data revealed that starch increase mainly took place in plates surrounding the pyrenoid during the day (1.4 fold and 9-fold increase between morning and afternoon in free-living and symbiotic cells, respectively). By contrast, the relative occupancy of starch grains in the cell did not vary between morning and afternoon in free-living and symbiotic cells (Fig. 4C, Table S9-10). The fact that transitory starch grains do not accumulate in symbiotic microalgae during the day suggests that starch degradation and sugar export into the cytosol could be maintained, like in actively growing free-living cells.

In order to further understand storage dynamics in free-living and symbiotic microalgae, starch quantification was also performed on the following day (after one night in the dark). We found that overnight consumption of starch was maintained in symbiotic microalgae despite their lower growth rate (similar diel starch turnover). In free-living (exponential growth phase) and symbiotic microalgae, about 44% of starch produced during the day was consumed overnight (from 1.40 ± 0.08 pg.cell^-1^ in the afternoon to 0.79 ± 0.02 pg.cell^-1^ the next morning in symbiotic microalgae) (Fig. 4A). Of note, this represents a 4.5-fold higher quantity of starch that is consumed in symbiosis overnight (0.61 pg.cell^-1^ *vs* 0.14 pg.cell^-1^ in free-living). This could indicate that nearly half of these carbohydrates/sugars have been i) tapped for algal energetic demands, ii) reallocated to other algal compartments (e.g. lipids), and/or iii) transferred to the host overnight in symbiosis.

### Accumulation of lipid droplets overnight in symbiotic microalgae

Our 24h-incubation nanoSIMS results showed that some of the fixed carbon can be used by symbiotic microalgae for their growth (^13^C enrichment in the algal cytoplasm), but also allocated into their cytosolic lipid droplets (Fig. 3B). It is known that lipid droplets are a major carbon storage compartment in microalgae that can fluctuate according to growth and stress conditions, such as nutrient limitation (Jouhet et al. 2022; Kong et al. 2018; Ran et al. 2019). Like starch, we tracked and quantified the neutral lipids content of free-living and symbiotic microalgae using a combination of a fluorescence-based assay (Nile Red staining quantified by spectrometry) and FIB-SEM imaging. The specificity of neutral lipid staining was verified in our microalgae using confocal fluorescence microscopy (Fig. S4). FIB-SEM-based 3D reconstruction revealed that symbiotic microalgae contained many more lipid droplets (up to 20) compared to the free-living stage (3 on average) (Fig. 5A). On average, the volume of total lipid droplets in symbiotic microalgae could be ∼20 times higher (or 11.7 times higher if normalized per cell volume: 0.035 % ± 0.007 µm^3^ in symbiosis compared to 0.003 % ± 0.003 µm^3^ in free-living cells) (Fig. 5B, Table S10. When compared between morning and afternoon, FIB-SEM and fluorescence-based quantification showed that there was no increase in lipid droplets in both free-living and symbiotic microalgae (Fig. 5). On the following day, lipid droplets did not increase in free-living cells, whereas in symbiosis, there was an accumulation of lipid droplets (0.019 ± 0.003 to 0.034 ± 0.009 a.u.cell^-1^) (Fig. 5C-Table S11). This accumulation in symbiotic microalgae seemed to be maintained the following days, up to day 7 (higher amount of lipid per cell). Overall, our results suggest that the storage of triacylglycerols (TAG) in lipids droplets increases overnight in symbiotic microalgae, likely fueled by carbon from starch, and that lipid droplets accumulate over consecutive days. This raises the question as to whether some of the photosynthetically-produced organic carbon is also transferred to the host.

**Figure 5.**
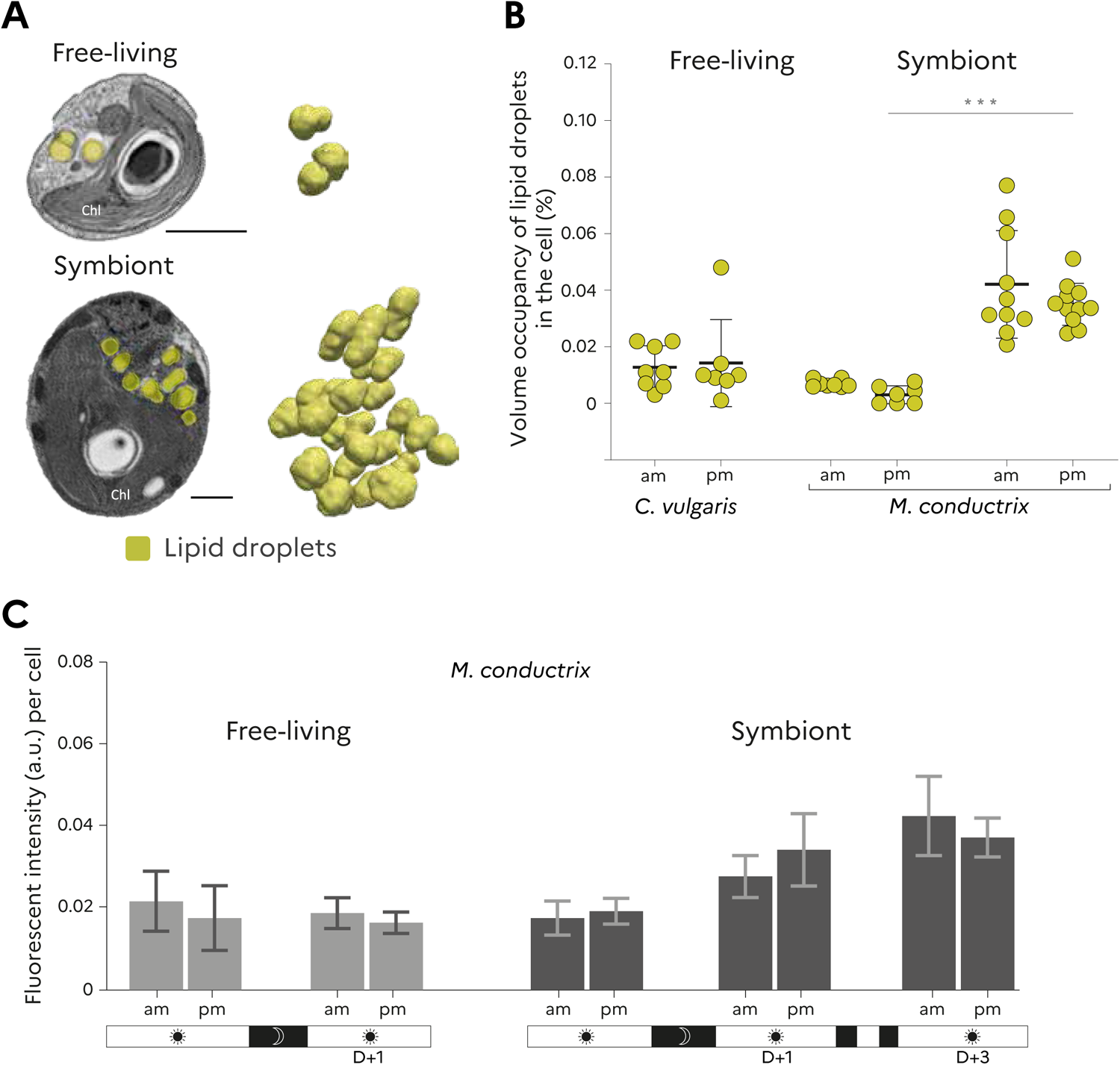
Accumulation of lipid droplets in symbiotic microalgae. **A)** Electron micrographs and FIB-SEM-based 3D reconstruction of lipid droplets (yellow) in free-living (*C. vulgaris* and *M. conductrix*) and symbiotic (*M. conductrix*) microalgae. Scale bar: 1 µm. **B)** FIB-SEM-based calculation of the volume of lipid droplets normalized by cell volume (occupancy in the algal cell as %) between free-living and symbiotic microalgae, and morning and afternoon. Statistical test: Non-parametric ANOVA unpaired test: ***p<0.0001: ns, no significant difference. **C)** Total neutral lipids per algal cell assessed using Nile Red staining in free-living and symbiotic microalgae in the morning and afternoon, over two consecutive days. In symbiotic microalgae, neutral lipids were also quantified after 7 days of culture. Fluorescence intensity (a.u.) was quantified by TECAN-based spectrometry. See also Supplementary Table S11 and Fig S4.

### Photosynthetically-derived carbon is transferred to the host

Using NanoSIMS, we investigated whether photosynthetically-derived carbon is transferred and stored in host cells. After 1h, no ^13^C enrichment was detected in the host cell (Figs. 3A and 6A). After 24h, we did not observe large structures/compartments of the host highly labelled with ^13^C, contrary to results reported for photosymbioses involving Foraminifera and corals (Gibbin et al. 2020; Kopp et al. 2015; Krueger et al. 2018; LeKieffre et al. 2018) that store carbon in large lipid droplets highly labelled with ^13^C. A low level of ^13^C enrichment was, however, detected in unknown host, demonstrating transfer to the host (Fig. 6A). We hypothesize that carbon energy produced by symbionts could be rapidly used by the host upon transfer and not stored in lipid droplets or other sugar reserves. To support this, we investigated the ultrastructural microenvironment of the host in the vicinity of symbionts. 3D reconstruction revealed a high density of host mitochondria surrounding symbiotic microalgae (Figs. 6B and 6C). Tight physical interaction between host mitochondria and the symbiont-containing symbiosome was also previously observed in this model (Song, Murata, and Suzaki 2017). Proximity between symbiotic microalgae and host mitochondria were also reported in salamander embryos and cnidarian-*Symbiodiniacae* symbioses (Dunn et al. 2012; Kerney et al. 2011). Therefore, it is possible that photosynthetically-derived organic carbon of symbionts could be transferred and very rapidly respired by host mitochondria, rendering it undetectable by nanoSIMS. This mass spectrometry imaging coupled with resin embedding can only detect carbon that is incorporated into biomass or stored in large molecules such as lipids and starch (Gibbin et al. 2020). We also cannot exclude that host mitochondria can participate to the delivery of CO_2_ surrounding symbiotic microalgae, so contributing to the enhanced algal carbon fixation and photosynthetic production.

**Figure 6.**
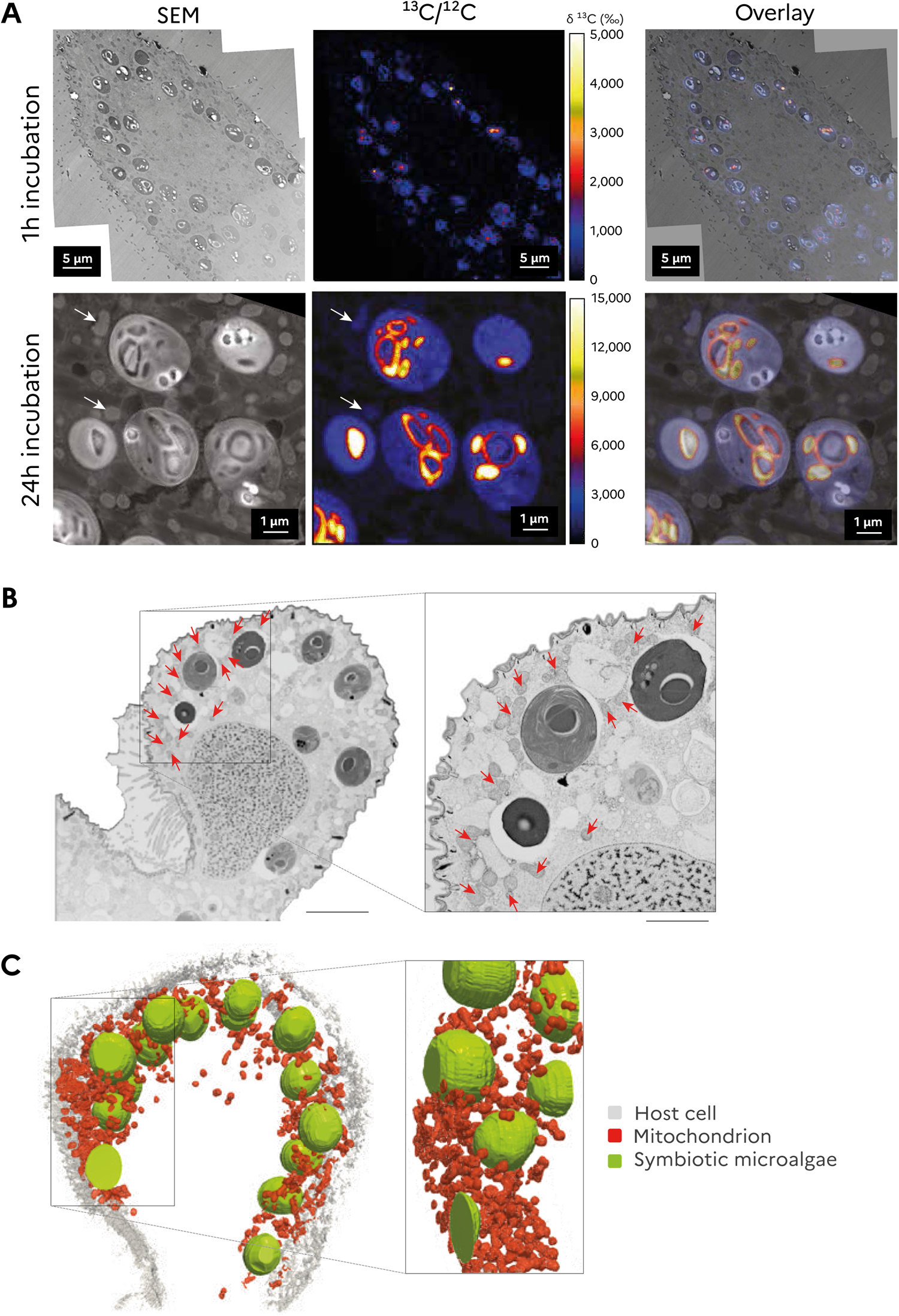
Carbon transfer in the host and aggregation of host mitochondria surrounding the symbiotic microalgae. **A)** Correlated SEM (Scanning Electron Microscope) and NanoSIMS images of the distribution of ^13^C enrichment in symbiotic microalgae of the host *Paramecium bursaria* after 1h and 24h-incubation with ^13^C-labelled bicarbonate. Colors in NanoSIMS maps represent enrichment relative to an unlabeled sample. At 24h incubation, white arrows indicate carbon transfer into unknown host structures. **B)** Electron micrograph from a FIB-SEM stack obtained from the host *P. bursaria* and its symbiotic microalgae. Red arrows indicate the host mitochondria. Scale bar: 1µm. **C)** 3D reconstruction of the host mitochondria (red) surrounding the symbiotic microalgae (green).

### Does the host act as a sink influencing carbon metabolism of its microalgae?

In plants and microalgae, photosynthesis and primary production are mainly driven by inputs such as light and CO_2_, but also by the balance between production and consumption of energy (Demmig-Adams et al. 2017; Krapp and Stitt 1995). This is the source-sink relationship, whereby production by the source (e.g. a microalga) can be enhanced by sinks (e.g. consumption for growth and/or export out of the site of production) (Abramson et al. 2016). Here, we show that carbon uptake and starch production in symbiotic microalgae are higher than in free-living cells while cell growth is lower. In addition, NanoSIMS results demonstrate transfer of some of this carbon to the host. We therefore hypothesize that the host could act as an additional sink, whereby its energetic demands influence photosynthetic production of its intracellular microalgae (source). This source-sink concept has also been proposed to be central in other symbiotic systems, from reefs to plants (Adams et al. 2020; Andersen 2003). To further understand the source-sink relationship in photosymbiosis, we quantified starch production of symbiotic microalgae when external glucose, considered to be one of the main photosynthates transferred (Arriola et al. 2018; Fujishima 2009; Sørensen et al. 2020), was provided to the host. An incubation experiment with ^13^C-glucose showed that the host *Paramecium bursaria* is able to take up this sugar molecule (9565.51 ‰ enrichment) (Table S12). We then compared starch production of symbiotic microalgae in a glucose-fed host and control (host without glucose) during six hours of light (from 10 am to 4 pm). This experiment revealed that symbiotic microalgae produced 2.3 times less starch at the end of the day when the host was provided with glucose (4.27 ± 0.73 pg.cell^-1^ in control *vs* 2.52 ± 0.36 pg.cell^-1^) (Fig. 7A). This lower starch production was not accompanied by a change in photosynthetic activity (net oxygen production) that remained similar in both conditions (Fig. S5, Table S13). Therefore, lower energetic demands of the host led to a cellular process that diminished starch production of its microalgae (but not light reactions of photosynthesis).

**Figure 7.**
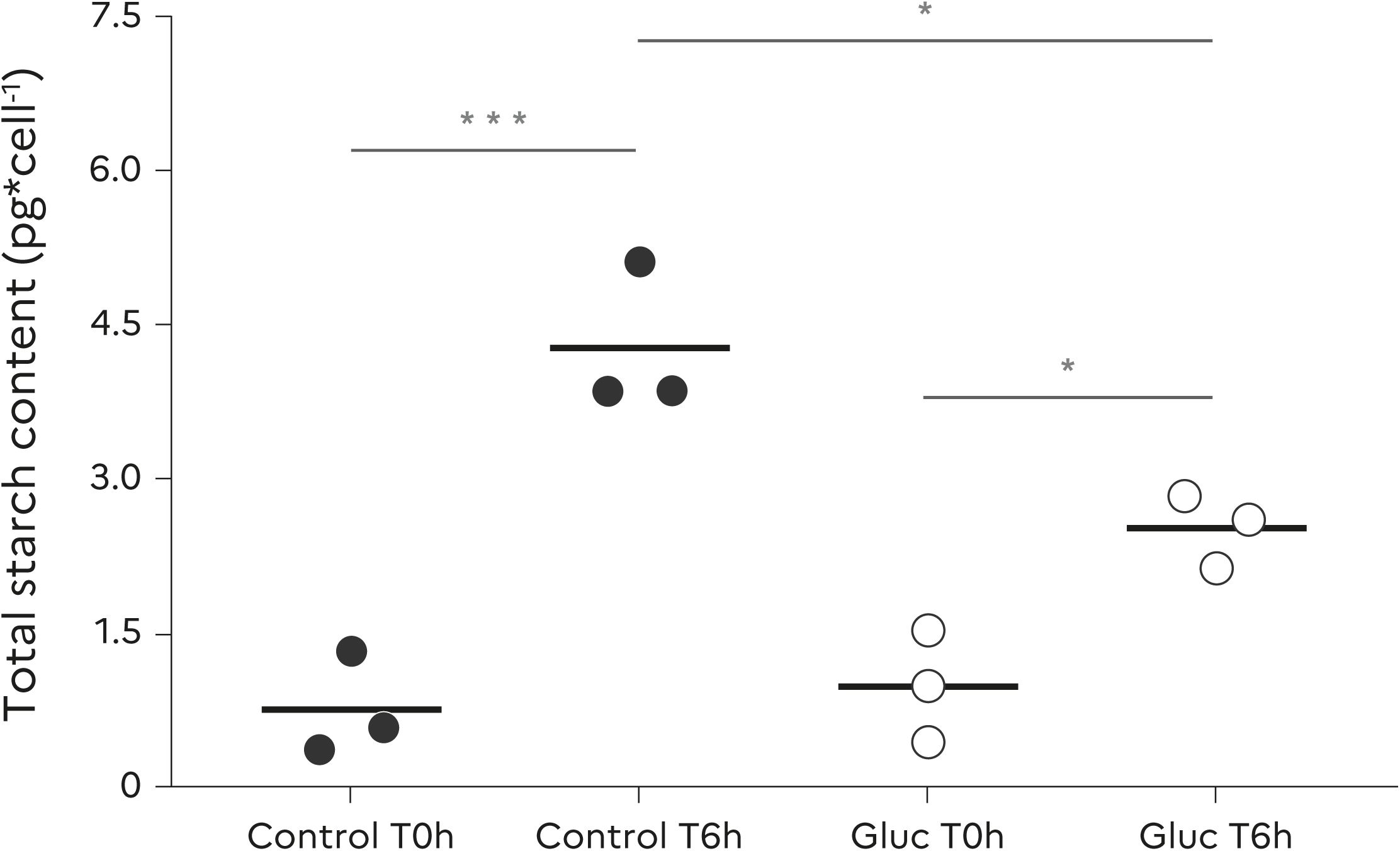
Starch production in symbiotic microalgae within hosts exposed to glucose. Total starch quantification of symbiont microalgae after exposure to glucose (75 mM) versus control (without glucose). Scatter plots shows the mean of biological triplicates± SD. ***p=0.0001: *p < 0.05. See also Supplementary Table S13 and Fig S5. (ANOVA, N = 12, Fcondition=5.8, Ftime=63.66; F interaction = 9.726).

## Conclusion and Perspectives

This multi-scale study provides clear evidence that photosynthetic production is enhanced in symbiotic microalgae compared to their free-living stage. We showed that symbiotic microalgae, the growth rate of which is repressed, have a 6-fold larger chloroplast with a 9-fold larger pyrenoid that contains 13 times more Rubisco. This is accompanied by 16-fold higher carbon fixation per microalgal cell. Enhancement of photosynthetic production in symbiotic microalgae therefore occurs in both marine (Uwizeye et al. 2021) and freshwater photosymbioses, suggesting common mechanisms within hosts. To date, the fate of this photosynthetically-derived carbon energy in photosymbiotic systems had not been fully addressed. Here, we demonstrate that the dynamics of diel starch turnover of the microalga is maintained in symbiosis, suggesting that the endogenous circadian clock known to regulate starch is maintained within a host (Graf and Smith 2011). However, symbiotic microalgae store more carbon as starch and neutral lipids compared to the free-living stage. More specifically, there is higher starch production during the day and higher consumption overnight, while neutral lipids (TAG) increase overnight and accumulate over successive days. Given the lower cell growth in symbiosis, these results indicate that symbiotic microalgae produce more organic carbon that needed for their growth. This excess of carbon energy stored in lipid droplets, which makes symbionts “fatty”, could nutritionally benefit the host when algal digestion takes place, a known phenomenon when the host is under starvation (Kodama and Miyazaki 2021). On a shorter time-scale, we also showed that the host can benefit from carbon exported by its microalgae, likely for sustaining its respiration needs, consistent with high density of host mitochondria surrounding the symbionts. Therefore, the host can act as an additional sink, likely influencing photosynthetic production by its microalgal symbionts, as indicated by the observation that symbiotic microalgae produce less starch when host energetic demands are lower. Nevertheless, lipid accumulation in symbiotic microalgae within the host *Paramecium bursaria* tends to show that the host is not a strong sink, in which case a massive import of photosynthetically-produced carbon would be observed. The host may potentially regulate carbon import from its microalgae based on its energetic demands in order to avoid uncontrolled efflux of carbohydrates that could be harmful for the system. Further studies are needed to fully investigate this source-sink relationship in photosymbiosis, which may be a foundational mechanistic process underlying the metabolic integration of microalgae and host cells.

## Material and Methods

### Strains and culture conditions

The ciliate *Paramecium bursaria* (CCAP1660/18) in symbiosis with *Micractinium conductrix* was obtained from the Culture Collection of Algae and Protozoa (https://www.ccap.ac.uk/). The culture medium for *P. bursaria* was prepared by inoculating Volvic natural mineral water with the bacterial strain *Serratia marcescens* CIP103235TI (Pasteur institute Bacteria CIP) and 0.66 g/L protozoan pellets (Carolina Biological Supply, NC, USA) 24h before use. The microalga *Chlorella vulgaris* (CCAP 211/11B) was also obtained from the Culture Collection of Algae and Protozoa. The *Micractinium conductrix culture* was obtained in culture by isolating its symbiotic stage within the ciliate host *P. bursaria* (CCAP1660/18). To do so, host cells were mechanically disrupted with a tissue grinder. Released symbiotic microalgae were recovered after filtration through a 40 µm cell strainer and 10 µm filter that removed host debris. The filtrate was centrifuged (2 min at 2 500 g) and plated on modified solid High Salt Medium (HSM) (Gorman and Levine 1965; Sueoka 1960). Microalgae were maintained at 20°C with a 12:12 h light/dark cycle - light intensity of 40 μE m^−2^ s^−1^ - and re-streaked on plates every week. After a period of growth, an individual colony of *M. conductrix* was re-streaked onto a fresh plate to establish a pure strain. All cultures (host and free-living microalgae) were maintained at 20°C under with a 12:12 h light/dark cycle with a light intensity of 40 μE m^−2^ s^−1^. Prior to carrying out experiments, free-living microalgal strains were transferred to liquid medium and maintained in the same conditions under constant agitation (80 rpm). The concentration of free-living and symbiotic microalgal cells was assessed with a LUNA-FLTM automated fluorescence cell counter (Logos Biosystems Inc., Anyang-si, Gyeonggi-do, South Korea).

### Physiological measurements: starch and neutral lipid quantification and photosynthetic oxygen measurements

Starch extraction was carried out by physicochemical disruption of cells following previous protocols (Wong et al. 2019). Briefly, cell cultures (triplicate samples for each experimental condition) were suspended in 1.2 ml of NaOH (1 M) in a centrifuge tube. The suspension was placed in a water bath at 90 °C for 10 min and then cooled to room temperature. Total starch content was then quantified with a commercial starch kit (Amylase/Amyloglucosidase Method - Product Code STA-20, SIGMA) and a spectrometer following the manufacturer’s instructions. The effect of external glucose on the total starch content of symbiotic microalgae was addressed by adding 75mM of glucose to the culture medium and incubating host cells containing symbiotic microalgae for 6 hours (from 10 am to 4 pm). Starch extraction and total starch quantification was carried out as described above. The control condition (without glucose) followed the same procedure. Total starch quantification is summarized in Table S7 and S13.

Neutral lipids was quantified by Nile Red (Sigma Aldrich) fluorescent staining (excitation wavelength at 532 nm and emission at 565 nm), as previously described (Abida et al. 2015; Cooksey et al. 1987). Cultures were first adjusted to a density of 1 million cells per ml. Nile Red solution (40 µl of a 2.5 µg.ml-1 stock solution in DMSO) was added to replicate 160 µl sub-samples of cell suspension in a black 96-well plate. Fluorescence was then measured using a TECAN infinite M1000 PRO (λ_ex_ = 530 nm). Nile Red staining was verified on our cells using confocal fluorescence microscope (Fig. S4). Micrographs showing the lipid droplets in microalgae were obtained using a Zeiss LSM 900 microscope using a 450–490-nm excitation filter. The results of neutral lipid quantification are summarized in Table S11.

Oxygen measurements was conducted following (Yee et al. 2023). Briefly, 500 µl of sample was used to measure oxygen in a WALZ KS-2500 water-jacketed chamber (Heinz Walz GmbH) paired with a FSO2-1 oxygen meter and optical microsensor (PyroScience GmbH). Samples were illuminated at 300 µmol photons m^-2^ sec^-1^ with stirring at 20° C in a MINI-PAM-II controlled by WinControl-3 software (Heinz Walz GmbH). Gross maximum oxygen production was calculated by the equation: O_2gross_ = O_2net_ – respiration. The results of O_2_ production rate analyses are summarized in Table S14.

### Sample preparation for electron microscopy

High-pressure freezing (HPM100, Leica) followed by freeze substitution (EM ASF2, Leica) was conducted to prepare samples for electron microscopy following the protocols of (Decelle et al 2019, 2021, Uwizeye et al 2021). Both free-living and symbiotic microalgae were harvested at exponential growth phase and concentrated by 2 min centrifugation at 2 500 g prior to cryo-fixation.

### Focused Ion Beam-Scanning Electron Microscopy (FIB-SEM)

Focused ion beam (FIB) tomography was performed with either a Zeiss NVision 40 or a Zeiss CrossBeam 550 microscope (Zeiss, Germany). The resin block containing the cells was fixed on a stub with carbon paste and surface-abraded with a diamond knife in a microtome to obtain a flat and clean surface. Samples were then metallized with 4 nm of platinum to avoid charging during observations. Inside the FIB-SEM, a second platinum layer (1–2 μm) was deposited locally on the analyzed area to mitigate possible curtaining artefacts. The sample was then abraded slice by slice with the Ga+ ion beam (generally with a current of 700 pA at 30 kV). Each exposed surface was imaged by scanning electron microscopy (SEM) at 1.5 kV and with a current of ∼1 nA using the in-lens EsB backscatter detector. In general, similar milling and imaging mode were used for all samples. Automatic correction of focus and astigmatism was performed during image acquisition, usually at approximately hourly intervals. For each slice, a thickness of 6 to 8 nm was removed, and SEM images were generally recorded with a pixel size between 6 to 8 nm, providing an isotropic voxel size. Whole volumes were imaged with 800–2000 frames, depending on the cell type and volume. Raw electron microscopy data are deposited in the Electron Microscopy Public Image Archive (EMPIAR), accession code EMPIAR-XXX.

### FIB-SEM analysis, segmentation and morphometrics

Image processing was initiated using software Fiji (https://imagej.net/Fiji) to crop selected cells and perform registration. Segmentation was based on pixel classification by a semi-automatic method adopted from (Uwizeye et al. 2020) using 3D slicer software (https://www.slicer.org/) and a supervised semi-automatic pixel classification mode (3 to 15 slices automatically segmented for each region of interest-ROI). Along with the cell, the main organelles and structures of the algal cells (nucleus, chloroplast, mitochondria, pyrenoid, starch and lipid droplets) were segmented. Morphometric analyses were calculated using the Statistics Module in 3D slicer. Results are provided in supplementary Tables S1-S4, S9-10.

### 13C bulk enrichment (Elemental Analyzer–Isotope Ratio Mass Spectrometry) and isotope analysis

Elemental analyzer isotope ratio mass spectrometry (EA-IRMS) and isotope analysis was conducted on free-living (*Chlorella vulgaris* and *Micractinium conductrix*) and symbiotic microalgae (*M. conductrix* from *P. bursaria* CCAP1660/18) to detect ^13^C-bicarbonate assimilation after 1 hour of treatment. An equivalent of 0.4 mg fresh weight (corresponding to 10^7^ cells) was used for the bulk analysis (Kimball et al. 1959). Cells were harvested at exponential growth phase (e.g. after 4 days of culture). For ^13^C enrichment, 10% of H^13^CO_3_ as a final concentration was used as the isotopic solution and added to modified HSM medium and bacterized Volvic-Pellet medium for free-living and symbiotic microalgae, respectively. After incubation, free-living algae were counted before centrifugation (15,000 rcf for 1 min at 20°C). Supernatant was discarded and the pellet was rinsed by three serial centrifugations, once with modified HSM and twice with MiliQ water. After the final centrifugation (15,000 rcf for 1 min at 20°C), cells were transferred into tin capsules for EA-IRMS analysis. For isolating symbiotic microalgae, host cells were mechanically disrupted by sonication (amplitude 40% * 2min, E 1J/s, 20ms - On/80ms - Off, Branson sonifier250) followed by a centrifugation (5 000 g for 2 min at 20°C) and two sequential filtration steps (40 µm cell strainer and 10 µm filter). The supernatant was discarded and symbiotic cells transferred into tin capsules for EA-IRMS analysis. All tin capsules were dried at room-temperature for one week before EA-IRMS analysis (LIENSs platform, La Rochelle). Control samples (unlabeled) were not incubated with ^13^C-labelled bicarbonate isotopes but otherwise followed the same steps. Carbon assimilation was calculated as described in (Uwizeye 2021). In brief, ^13^C-uptake per cell in free-living and symbiotic microalgae was estimated from the calculated ^13^C-excess and averaged total carbon. Data was expressed as carbon uptake per cell and carbon uptake normalized per carbon (Table S5). In the same experimental conditions, we also incubated cells with ^13^C-labelled glucose for 24h to see whether host cells can uptake this molecule. 2.5 mL of ^13^C-glucose was added in the culture flask to reach a final concentration of 1mM. Results are provided in Table S12.

### Rubisco quantification

Total protein extracts were obtained from microalgal cells in exponential phase of growth (e.g. 4 days) in 50 mM Tris buffer pH 8.0 supplemented with protein inhibitor cocktail (539131, Calbiochem). Free-living and symbiotic microalgae were disrupted by bead beating with a Precellys device (Bertin Technologies) using micro glass beads (500 µm) with two 30 seconds cycles at 5000 rpm. After centrifugation, proteins in the supernatant were precipitated overnight at −20°C in 100 % acetone. After a second centrifugation, the pellet was solubilized for 5 min (RT) in 50 mM Tris (pH 6.8), 2% sodium dodecyl sulphate, 10 mM EDTA, and protein inhibitor cocktail. After a second centrifugation, supernatant was retained and protein quantified with the DC Protein assay kit II (Biorad). Proteins samples (1.5µg and 3µg of proteins) of free living (*C. vulgaris and M. conductrix*) and symbiotic microalgae (*M. conductrix* from *P. bursaria* CCAP1660/18) were loaded on 10% SDS-PAGE gels (Mini-PROTEAN TGX Precast Protein Gels, Biorad) and blotted onto nitrocellulose membranes. A Rubisco positive control (AS01017S, Agrisera) was used to generate a standard curve. Membranes were blocked for 1h with 5% low fat milk powder in TBS-T Tween 0.1% and probed with anti-RBCL antibody (AS03037, Agrisera, 1:10000, ON) and secondary HRP conjugated anti rabbit antibody (111-035-003) (Interchim, 1:10000, 1h) in TBS-T containing 5% low fat milk powder. Antibody incubations were followed by washing in TBS-T. All steps were performed at room temperature with agitation. Blots were developed for 1min with ECL Prime detection kit (RPN2232, Amersham) according the manufacturer’s instructions (GE Heathcare). Images of the blot were obtained using a CCD imager (Chemidoc MP system, Biorad) and ImageJ software. Data was expressed as rubisco per cell (Table S6).

## Data analysis

Morphometric data was analyzed with Graphpad Prism 6 software and R studio. Statistical comparisons were performed with non-parametric unpaired ANOVA for a multiple comparison with Dunn’s test correction.

## Data Availability

Raw 3D electron microscopy images data have been deposited on EMPIAR: DOI:. All other study data are included in the article and/or supporting information.

## Acknowledgments

This project received funding from the LabEx GRAL (ANR-10-LABX-49-01), financed within the University Grenoble Alpes graduate school (Ecoles Universitaires de Recherche) CBHEUR-GS (ANR-17-EURE-0003). JD was supported by CNRS and ATIP-Avenir program funding. We thank Anders Meibom and Stephane Escrig for help with NanoSIMS analysis. We also thank the EA-IRMS platform (Gael Guillou and Benoit Lebreton) - a FEDER funding. We thank Guy Schoehn and Christine Moriscot, and the electron microscope facility at IBS, which is supported by the Rhône-Alpes Region, the Fondation Recherche Medicale (FRM), the fonds FEDER, the Center National de la Recherche Scientifique (CNRS), the CEA, the University of Grenoble, EMBL, and the GIS Infrastructures en Biologie Sante et Agronomie (IBISA). We thank Ian Probert for critically reading the manuscript and suggesting improvements.

## Author contributions

A.C.G. and J.D. designed research, interpreted results, and drafted the manuscript with the support of G.C. J.D. supervised the work. A.C.G., B.G. and J.D. jointly performed the sample preparation for microscopy. P.H.J., N.L.S. and Y.S. conducted FIB-SEM imaging, and A.C.G. analyzed the data. C.L. conducted nanoSIMS analyses and analyzed the data. F.C. and D.Y. performed and assisted with experiments. A.C.G. and J.D. wrote the manuscript with G.C.

