## Supplementary materials for "Sweet and fatty symbionts: photosynthetic productivity and carbon storage boosted in microalgae within a host"

*Corresponding author: Johan Decelle and Andrea Catacora-Grundy

This PDF file includes: Figs. S1 to S5 – Table S1-S13

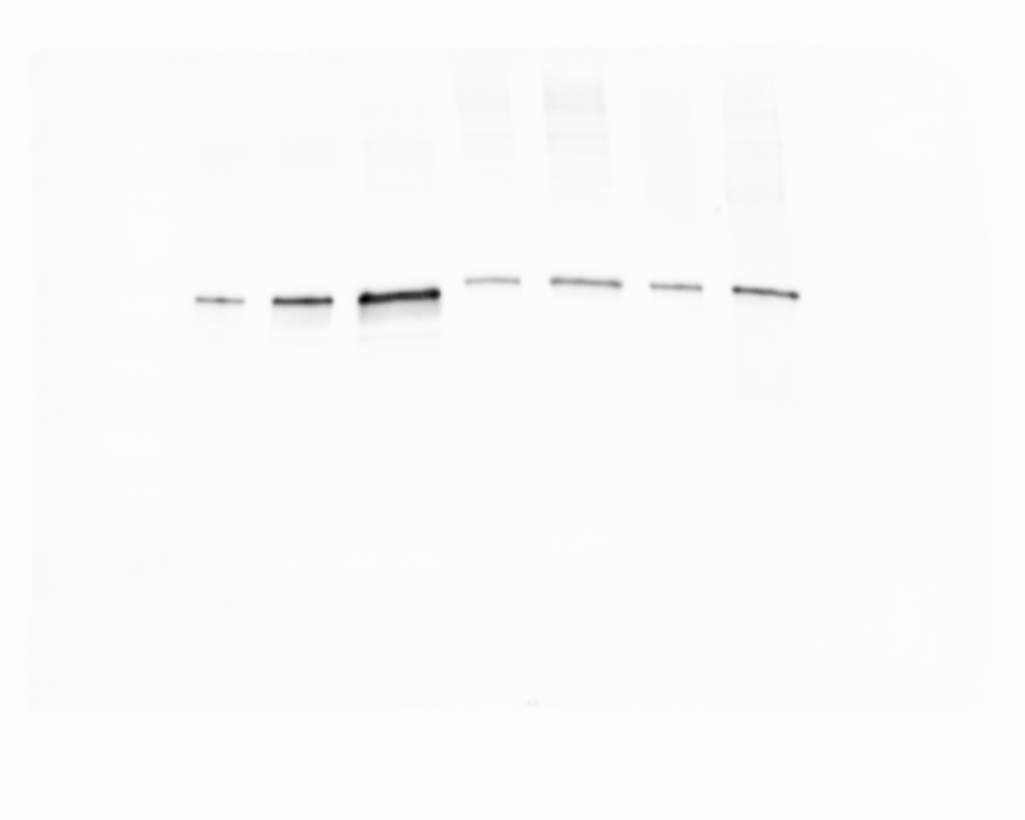

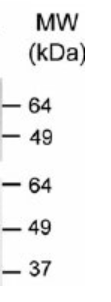

Rubisco

Standard

Free-living

Symbiont

0.3

0.75

1.5

1.5

1.5

3

3

[p.mol]

**Fig. S1**. Western blot analysis for Ribulose-1,5-bisphosphate carboxylase (RuBisCo) quantification. First 3 lanes represent different concentrations (picomole) of Rubisco as standard. The remaining lanes correspond to Rubisco protein extracted from free-living (*M. conductrix;* 2.3*10^8^ total cells) and symbiotic microalgae (*M. conductrix* in symbiosis with *P. bursaria* CCAP1660/18; 2*10^7^ total cells) in two different quantities (1.5 µg and 3 µg).

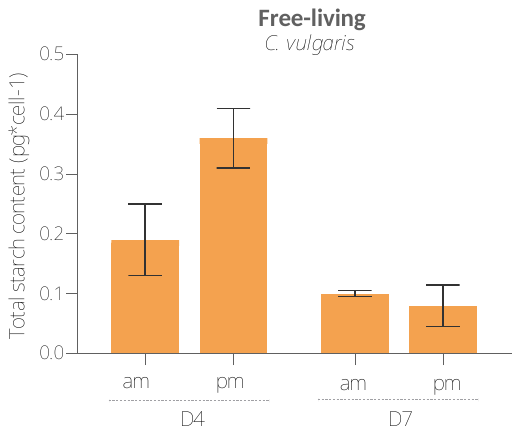

**Fig. S2.** Total starch quantification of the free-living *Chlorella vulgaris.* Starch quantification was assessed by enzymatic assay (Amylase/Amyloglucosidase Method - Product Code STA-20, SIGMA) morning and afternoon at day 4=D4 and day 7=D7 of culture, corresponding to the mid and late exponential growth phase. Bar plots represent the mean of total starch content per cell of biological triplicates ± SE.

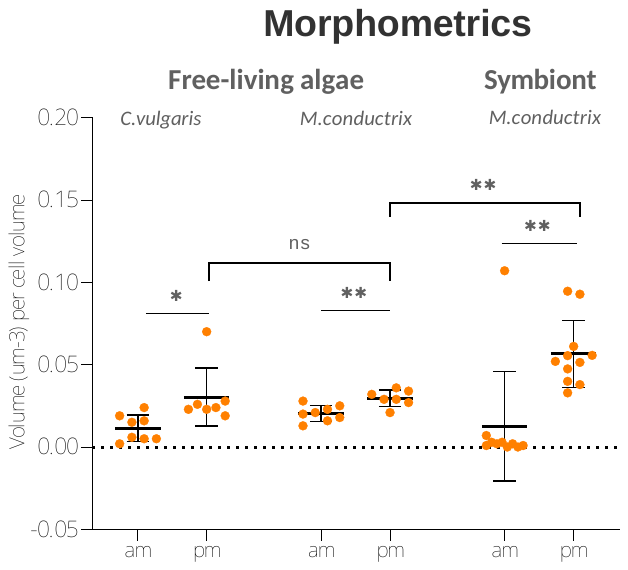

**Fig. S3.** Volume of total starch per algal cell volume (% occupancy) calculated from FIB-SEM volumetrics between both free-living microalgae (*C. vulgaris* and *M. conductrix*) and symbiotic microalgae (*M. conductrix* of *P. bursaria* CCAP1660/18) harvested in the morning and afternoon.

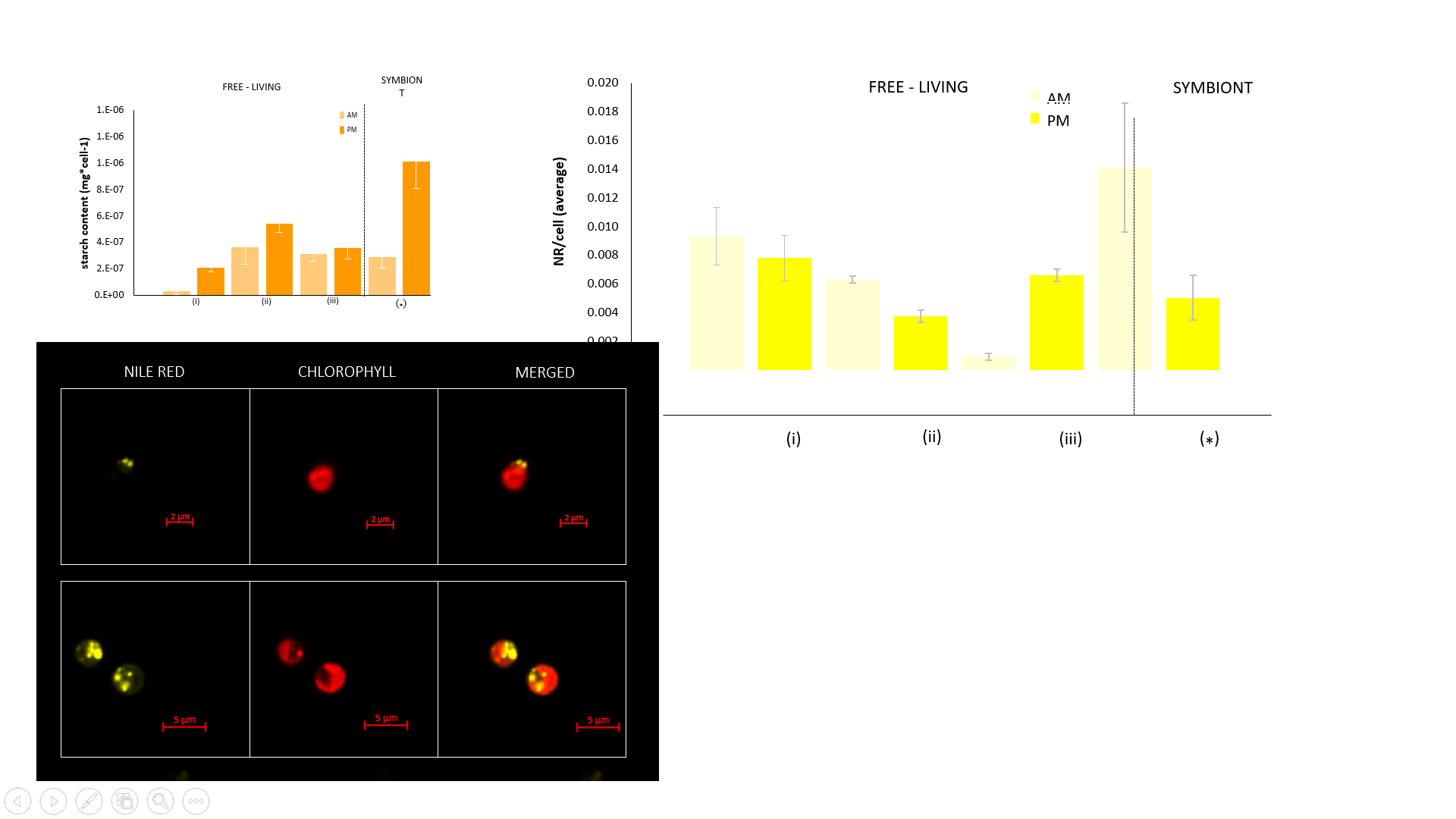

**Fig. S4.** Nile Red staining coupled with confocal fluorescence microscopy to observe neutral lipids in free-living (*M. conductrix*) and symbiotic microalgae (*M. conductrix* of *P. bursaria* CCAP1660/18). Nile red staining shows that neutral lipids are mainly localized in lipid droplets (yellow) in the algal cytoplasm, outside the chloroplast (visible in red by chlorophyll autofluorescence).

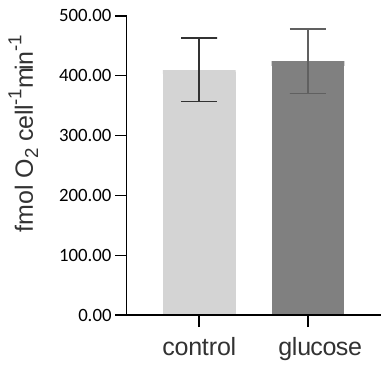

**Fig. S5.** Measurements of gross maximum photosynthetic oxygen production of *Paramecium bursaria* in symbiosis with *Micractinium conductrix* exposed glucose (75 mM) compared to the control for 6 hours in light). Bar plots show the mean values of triplicate measurements and ± SEM.

| ***Micractinium conductrix*** | Cell_1_ (*µm^3^*) | Cell_2_ (*µm^3^*) | Cell_3_ (*µm^3^*) | Cell_4_ (*µm^3^*) | Cell_5_ (*µm^3^*) | Cell_6_ (*µm^3^*) | Cell_7_ (*µm^3^*) | Cell_8_ (*µm^3^*) | Cell_9_ (*µm^3^*) | Cell_10_ (*µm^3^*) | **Average (*µm^3^*)** | **STD (*µm^3^*)** |
| --- | --- | --- | --- | --- | --- | --- | --- | --- | --- | --- | --- | --- |
| total starch | 0.20 | 0.08 | 0.31 | 0.14 | 0.11 | 0.11 | 0.26 | 0.27 | 0.19 | 0.28 | 0.20 | 0.08 |
| starch plates | 0.20 | 0.08 | 0.30 | 0.11 | 0.24 | 0.26 | 0.14 | 0.11 | 0.18 | 0.27 | 0.19 | 0.08 |
| starch grains | 0.00 | 0.00 | 0.02 | 0.00 | 0.02 | 0.00 | 0.00 | 0.00 | 0.01 | 0.01 | 0.01 | 0.01 |
| lipid droplets | 0.07 | 0.06 | 0.07 | 0.04 | 0.07 | 0.07 | 0.04 | 0.06 | 0.05 | 0.04 | 0.06 | 0.02 |
| cell | 10.41 | 6.10 | 11.07 | 5.54 | 8.22 | 9.26 | 6.06 | 6.58 | 6.56 | 8.08 | 7.79 | 1.94 |
| pyrenoid | 0.17 | 0.09 | 0.20 | 0.09 | 0.16 | 0.11 | 0.09 | 0.10 | 0.11 | 0.18 | 0.13 | 0.04 |
| nucleus | 0.97 | 0.79 | 1.04 | 0.76 | 0.86 | 0.95 | 0.76 | 0.78 | 0.74 | 0.83 | 0.85 | 0.11 |
| mitochondrion | 0.29 | 0.16 | 0.31 | 0.18 | 0.22 | 0.22 | 0.17 | 0.23 | 0.17 | 0.19 | 0.21 | 0.05 |
| chloroplast | 5.12 | 2.79 | 5.20 | 2.32 | 3.83 | 4.41 | 2.69 | 3.09 | 2.97 | 3.49 | 3.59 | 1.02 |

| ***Micractinium conductrix* (isolated from *P. bursaria CCAP 1660/18)*** | Cell_1_ (*µm^3^*) | Cell_2_ (*µm^3^*) | Cell_3_ (*µm^3^*) | Cell_4_ (*µm^3^*) | Cell_5_ (*µm^3^*) | Cell_6_ (*µm^3^*) | Cell_7_ (*µm^3^*) | Cell_8_ (*µm^3^*) | Cell_9_ (*µm^3^*) | Cell_10_ (*µm^3^*) | **Average (*µm^3^*)** | **STD (*µm^3^*)** |
| --- | --- | --- | --- | --- | --- | --- | --- | --- | --- | --- | --- | --- |
| total starch | 2.90 | 3.17 | 2.37 | 2.12 | 1.72 | 2.11 | 2.24 | 0.34 | 0.40 | 0.28 | 1.77 | 1.07 |
| starch plates | 2.56 | 2.29 | 1.99 | 1.99 | 1.60 | 2.09 | 1.75 | 0.34 | 0.40 | 0.28 | 1.53 | 0.86 |
| starch grains 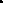 | 0.35 | 0.89 | 0.38 | 0.13 | 0.13 | 0.02 | 0.49 | 0.00 | 0.00 | 0.00 | 0.24 | 0.29 |
| lipid droplets | 0.91 | 1.33 | 0.80 | 0.92 | 0.64 | 0.68 | 1.00 | 0.63 | 0.45 | 0.76 | 0.81 | 0.25 |
| cell | 51.61 | 59.18 | 55.69 | 47.93 | 42.39 | 45.84 | 54.39 | 28.06 | 26.22 | 24.23 | 43.55 | 12.97 |
| pyrenoid | 1.06 | 1.44 | 1.39 | 1.35 | 1.29 | 0.93 | 1.39 | 1.10 | 0.83 | 0.65 | 1.14 | 0.27 |
| nucleus | 2.53 | 2.60 | 3.35 | 2.40 | 2.05 | 1.83 | 2.46 | 1.81 | 1.67 | 1.84 | 2.25 | 0.52 |
| mitochondrion | 1.76 | 1.94 | 2.31 | 1.67 | 1.45 | 1.88 | 1.86 | 1.01 | 0.82 | 0.78 | 1.55 | 0.52 |
| chloroplast | 27.60 | 32.14 | 30.07 | 26.28 | 23.72 | 24.04 | 29.38 | 12.07 | 12.76 | 11.18 | 22.92 | 7.97 |

| ***Chlorella vulgaris*** | Cell_1_ (*µm^3^*) | Cell_2_ (*µm^3^*) | Cell_4_ (*µm^3^*) | Cell_5_ (*µm^3^*) | Cell_8_ (*µm^3^*) | Cell_9_ (*µm^3^*) | **Average (*µm^3^*)** | **STD (*µm^3^*)** |
| --- | --- | --- | --- | --- | --- | --- | --- | --- |
| total starch | 0.14 | 0.36 | 0.01 | 0.07 | 0.39 | 2.12 | 0.51 | 0.80 |
| starch plates | 0.14 | 0.36 | 0.01 | 0.07 | 0.37 | 0.42 | 0.23 | 0.18 |
| starch grains | 0.00 | 0.00 | 0.00 | 0.00 | 0.02 | 1.70 | 0.29 | 0.69 |
| lipid droplets | 0.36 | 0.43 | 0.06 | 0.21 | 0.22 | 0.13 | 0.24 | 0.14 |
| cell | 10.27 | 13.82 | 5.49 | 10.26 | 15.89 | 17.49 | 12.20 | 4.40 |
| pyrenoid | 0.23 | 0.29 | 0.06 | 0.13 | 0.22 | 0.18 | 0.18 | 0.08 |
| nucleus | 0.65 | 0.93 | 0.70 | 1.07 | 1.53 | 0.75 | 0.94 | 0.33 |
| mitochondrion | 0.40 | 0.66 | 0.18 | 0.37 | 0.47 | 0.59 | 0.45 | 0.17 |
| chloroplast | 3.70 | 5.00 | 2.40 | 4.27 | 7.55 | 6.93 | 4.98 | 1.96 |

**Table S1.** FIB-SEM based morphometrics of free-living (*Chlorella vulgaris* and *Micractinium conductrix*) and symbiotic (*M. conductrix* of *P. bursaria* CCAP1660/18) microalgae revealing the volume of the cell, total starch, lipid droplets, pyrenoid, nucleus, mitochondrion and chloroplast (µm^3^).

| ***Micractinium conductrix*** | Cell_1_ (%) | Cell_2_ (%) | Cell_3_ (%) | Cell_4_ (%) | Cell_5_ (%) | Cell_6_ (%) | Cell_7_ (%) | Cell_8_ (%) | Cell_9_ (%) | Cell_10_ (%) | **Average (%)** | **STD (%)** |
| --- | --- | --- | --- | --- | --- | --- | --- | --- | --- | --- | --- | --- |
| starch plates | 1.97 | 1.32 | 2.67 | 2.26 | 1.65 | 2.05 | 2.94 | 2.84 | 2.72 | 3.29 | 2.37 | 0.62 |
| starch grains | 0.00 | 0.00 | 0.17 | 0.06 | 0.00 | 0.02 | 0.22 | 0.04 | 0.17 | 0.16 | 0.08 | 0.08 |
| lipid droplets | 0.70 | 0.90 | 0.59 | 0.58 | 0.90 | 0.68 | 0.91 | 0.77 | 0.69 | 0.49 | 0.72 | 0.15 |
| cell | 34.47 | 35.03 | 35.68 | 36.09 | 33.62 | 36.91 | 34.17 | 34.83 | 35.67 | 37.94 | 35.44 | 1.31 |
| pyrenoid | 1.63 | 1.53 | 1.78 | 1.49 | 1.50 | 1.56 | 1.97 | 1.23 | 1.61 | 2.25 | 1.66 | 0.29 |
| nucleus | 9.31 | 12.88 | 9.41 | 12.47 | 11.80 | 13.73 | 10.49 | 10.25 | 11.23 | 10.24 | 11.18 | 1.50 |
| mitochondrion | 2.77 | 2.59 | 2.76 | 2.76 | 3.51 | 3.17 | 2.72 | 2.37 | 2.61 | 2.41 | 2.77 | 0.34 |
| chloroplast | 49.16 | 45.76 | 46.94 | 44.28 | 47.02 | 41.86 | 46.58 | 47.67 | 45.31 | 43.22 | 45.78 | 2.18 |

| ***Micractinium conductrix (isolated from P. bursaria CCAP 1660/18)*** | Cell_1_ (%) | Cell_2_ (%) | Cell_3_ (%) | Cell_4_ (%) | Cell_5_ (%) | Cell_6_ (%) | Cell_7_ (%) | Cell_8_ (%) | Cell_9_ (%) | Cell_10_ (%) | **Average (%)** | **STD (%)** |
| --- | --- | --- | --- | --- | --- | --- | --- | --- | --- | --- | --- | --- |
| starch plates | 4.95 | 3.87 | 3.57 | 4.15 | 3.77 | 4.56 | 3.21 | 1.21 | 1.54 | 1.14 | 3.20 | 1.40 |
| starch grains | 0.67 | 1.50 | 0.68 | 0.27 | 0.30 | 0.04 | 0.90 | 0.00 | 0.00 | 0.00 | 0.44 | 0.50 |
| lipid droplets | 1.77 | 2.24 | 1.44 | 1.92 | 1.50 | 1.49 | 1.84 | 2.25 | 1.70 | 3.12 | 1.93 | 0.51 |
| cell | 28.75 | 27.99 | 27.66 | 27.53 | 27.18 | 31.37 | 29.54 | 39.55 | 35.43 | 36.11 | 31.11 | 4.38 |
| pyrenoid | 2.06 | 2.43 | 2.49 | 2.81 | 3.03 | 2.02 | 2.55 | 3.92 | 3.15 | 2.68 | 2.71 | 0.56 |
| nucleus | 4.91 | 4.39 | 6.01 | 5.01 | 4.84 | 3.99 | 4.52 | 6.45 | 6.39 | 7.59 | 5.41 | 1.14 |
| mitochondrion | 3.41 | 3.28 | 4.16 | 3.48 | 3.43 | 4.10 | 3.42 | 3.61 | 3.12 | 3.21 | 3.52 | 0.35 |
| chloroplast | 53.48 | 54.30 | 53.99 | 54.83 | 55.95 | 52.44 | 54.01 | 43.01 | 48.68 | 46.15 | 51.68 | 4.27 |

| ***Chlorella vulgaris*** |  | Cell_1_ (%) | Cell_2_ (%) | Cell_4_ (%) | Cell_5_ (%) | Cell_8_ (%) | Cell_9_ (%) | **Average (%)** | **STD (%)** |
| --- | --- | --- | --- | --- | --- | --- | --- | --- | --- |
| starch plates |  | 1.32 | 2.59 | 0.17 | 0.65 | 2.33 | 2.40 | 1.58 | 1.02 |
| starch grains |  | 0.00 | 0.00 | 0.00 | 0.00 | 0.12 | 9.73 | 1.64 | 3.96 |
| lipid droplets |  | 3.54 | 3.11 | 1.12 | 2.06 | 1.39 | 0.75 | 2.00 | 1.12 |
| cell |  | 46.70 | 44.53 | 37.86 | 40.30 | 34.74 | 38.82 | 40.49 | 4.42 |
| pyrenoid |  | 2.20 | 2.10 | 1.06 | 1.29 | 1.36 | 1.02 | 1.51 | 0.52 |
| nucleus |  | 6.29 | 6.74 | 12.67 | 10.43 | 9.61 | 4.27 | 8.34 | 3.10 |
| mitochondrion |  | 3.90 | 4.77 | 3.34 | 3.61 | 2.93 | 3.38 | 3.66 | 0.63 |
| chloroplast |  | 36.05 | 36.16 | 43.78 | 41.66 | 47.51 | 39.63 | 40.80 | 4.47 |

**Table S2.** Volume occupancy (organelle-volume/cell-volume ratio) based on FIB-SEM morphometrics in free-living (*Chlorella vulgaris* and *Micractinium conductrix*) and symbiotic (*M. conductrix* of *P. bursaria* CCAP1660/18) microalgae of starch, lipid droplets, pyrenoid, nucleus, mitochondrion and chloroplast (%).

| ***Micractinium conductrix*** | Cell_1_ (*µm^3^*) | Cell_2_ (*µm^3^*) | Cell_3_ (*µm^3^*) | Cell_4_ (*µm^3^*) | Cell_5_ (*µm^3^*) | Cell_6_ (*µm^3^*) | Cell_7_ (*µm^3^*) | Cell_8_ (*µm^3^*) | Cell_9_ (*µm^3^*) | Cell_10_ (*µm^3^*) | **Average (*µm^3^*)** | **STD (*µm^3^*)** |
| --- | --- | --- | --- | --- | --- | --- | --- | --- | --- | --- | --- | --- |
| cell | 10.41 | 6.10 | 11.07 | 6.06 | 6.58 | 5.54 | 8.22 | 9.26 | 6.56 | 8.08 | 7.79 | 1.94 |
| pyrenoid | 0.17 | 0.09 | 0.20 | 0.09 | 0.10 | 0.09 | 0.16 | 0.11 | 0.11 | 0.18 | 0.13 | 0.04 |
| chloroplast | 5.12 | 2.79 | 5.20 | 2.69 | 3.09 | 2.32 | 3.83 | 4.41 | 2.97 | 3.49 | 3.59 | 1.02 |
| ***Micractinium conductrix* (isolated from *P. bursaria CCAP 1660/18)*** | Cell_1_ (*µm^3^*) | Cell_2_ (*µm^3^*) | Cell_3_ (*µm^3^*) | Cell_4_ (*µm^3^*) | Cell_5_ (*µm^3^*) | Cell_6_ (*µm^3^*) | Cell_7_ (*µm^3^*) | Cell_8_ (*µm^3^*) | Cell_9_ (*µm^3^*) | Cell_10_ (*µm^3^*) | **Average (*µm^3^*)** | **STD (*µm^3^*)** |
| cell | 51.61 | 59.18 | 55.69 | 47.93 | 42.39 | 45.84 | 54.39 | 28.06 | 26.22 | 24.23 | 43.55 | 12.97 |
| pyrenoid | 1.06 | 1.44 | 1.39 | 1.35 | 1.29 | 0.93 | 1.39 | 1.10 | 0.83 | 0.65 | 1.14 | 0.27 |
| chloroplast | 27.60 | 32.14 | 30.07 | 26.28 | 23.72 | 24.04 | 29.38 | 12.07 | 12.76 | 11.18 | 22.92 | 7.97 |

| ***Chlorella vulgaris*** | Cell_1_ (*µm^3^*) | Cell_2_ (*µm^3^*) | Cell_4_ (*µm^3^*) | Cell_5_ (*µm^3^*) | Cell_8_ (*µm^3^*) | Cell_9_ (*µm^3^*) | **Average (*µm^3^*)** | **STD (*µm^3^*)** |
| --- | --- | --- | --- | --- | --- | --- | --- | --- |
| cell | 10.27 | 13.82 | 5.49 | 10.26 | 15.89 | 17.49 | 12.20 | 4.40 |
| pyrenoid | 0.23 | 0.29 | 0.06 | 0.13 | 0.22 | 0.18 | 0.18 | 0.08 |
| chloroplast | 3.70 | 5.00 | 2.40 | 4.27 | 7.55 | 6.93 | 4.98 | 1.96 |

**Table S3**. FIB-SEM based morphometrics of free-living (*Chlorella vulgaris* and *Micractinium conductrix*) and symbiotic (*M. conductrix* of *P. bursaria* CCAP1660/18) microalgae revealing the volume of the pyrenoid and chloroplast (µm^3^).

| ***Chlorella vulgaris*** | Cell_1_ (%) | Cell_2_ (%) | Cell_4_ (%) | Cell_5_ (%) | Cell_8_ (%) | Cell_9_ (%) | **Average (%)** | **STD (%)** |
| --- | --- | --- | --- | --- | --- | --- | --- | --- |
| pyrenoid | 6.11 | 5.80 | 2.42 | 3.10 | 2.87 | 2.57 | 3.81 | 1.68 |
| chloroplast | 93.89 | 94.20 | 97.58 | 96.90 | 97.13 | 97.43 | 96.19 | 1.68 |

| ***Micractinium conductrix*** | Cell_1_ (%) | Cell_2_ (%) | Cell_3_ (%) | Cell_4_ (%) | Cell_5_ (%) | Cell_6_ (%) | Cell_7_ (%) | Cell_8_ (%) | Cell_9_ (%) | Cell_10_ (%) | **Average (%)** | **STD (%)** |
| --- | --- | --- | --- | --- | --- | --- | --- | --- | --- | --- | --- | --- |
| pyrenoid | 3.31 | 3.34 | 3.80 | 3.37 | 3.19 | 3.74 | 4.23 | 2.59 | 3.55 | 5.22 | 3.63 | 0.70 |
| chloroplast | 96.69 | 96.66 | 96.20 | 96.63 | 96.81 | 96.26 | 95.77 | 97.41 | 96.45 | 94.78 | 96.37 | 0.70 |

| ***Micractinium conductrix* (isolated from *P. bursaria CCAP 1660/18)*** | Cell_1_ (%) | Cell_2_ (%) | Cell_3_ (%) | Cell_4_ (%) | Cell_5_ (%) | Cell_6_ (%) | Cell_7_ (%) | Cell_8_ (%) | Cell_9_ (%) | Cell_10_ (%) | **Average (%)** | **STD (%)** |
| --- | --- | --- | --- | --- | --- | --- | --- | --- | --- | --- | --- | --- |
| pyrenoid | 3.85 | 4.48 | 4.61 | 5.13 | 5.42 | 3.86 | 4.72 | 9.11 | 6.47 | 5.81 | 5.35 | 1.56 |
| chloroplast | 96.15 | 95.52 | 95.39 | 94.87 | 94.58 | 96.14 | 95.28 | 90.89 | 93.53 | 94.19 | 94.65 | 1.56 |

**Table S4.** Volume occupancy (organelle-volume/cell-volume ratio, %) of the pyrenoid within the chloroplast based on FIB-SEM morphometrics in free-living (*Chlorella vulgaris* and *Micractinium conductrix*) and symbiotic (*M. conductrix* of *P. bursaria* CCAP1660/18) microalgae.

| **Sample** | **Isotope** | **condition / time** | **Replicate** | **Number of cells** | **Iiso**  (ng ^13^C/sample) | **Iiso**  (pg ^13^C.cell^-1^) | **Iiso**  (pg ^13^C. pg C^-1^) |
| --- | --- | --- | --- | --- | --- | --- | --- |
| Free-living *M. conductrix* | ^13^C | 1h | 1 | 4.20E+07 | 42.03 | **0.0010** | **0.0008** |
| Free-living *M. conductrix* | ^13^C | 1h | 2 | 1.63E+08 | 203.51 | **0.0012** | **0.0013** |
| Free-living *M. conductrix* | ^13^C | 1h | 3 | 6.30E+07 | 88.47 | **0.0014** | **0.0012** |
|  | ^13^C |  |  |  |  |  |  |
| Symbiotic *M. conductrix* (1660/18) | ^13^C | 1h | 1 | 1.17E+07 | 391.84 | **0.0335** | **0.0024** |
| Symbiotic *M. conductrix* (1660/18) | ^13^C | 1h | 2 | 1.90E+07 | 204.96 | **0.0108** | **0.0023** |
| Symbiotic *M. conductrix* (1660/18) | ^13^C | 1h | 3 | 1.70E+07 | 224.15 | **0.0132** | **0.0020** |

| **Sample** | **Isotope** | **condition / time** | ** Iiso**  (ng ^13^C/sample) | ** Iiso**  (pg ^13^C.cell^-1^) | **SD** | ** Iiso**  (pg ^13^C. pg C^-1^) | **SD** |
| --- | --- | --- | --- | --- | --- | --- | --- |
| Free-living *M. conductrix* | ^13^C | 1h | 111.34 | **0.0012** | 0.0002 | **0.0011** | 0.0003 |
| Symbiotic *M. conductrix* (1660/18) | ^13^C | 1h | 273.65 | **0.0192** | 0.0125 | **0.0022** | 0.0002 |

**Table S5**. Carbon uptake rate calculated based on 1h incubation with ^13^C-bicarbonate in free-living (*Micractinium conductrix*) and symbiotic (*M. conductrix* of *P. bursaria* CCAP1660/18) microalgae.

|  | **Rubisco* pmol** | | | | | | **Rubisco**  (pmol/cell) | | | | | | | **Rubisco**  (pmol / µg prot / cell) | | | | | |
| --- | --- | --- | --- | --- | --- | --- | --- | --- | --- | --- | --- | --- | --- | --- | --- | --- | --- | --- | --- |
| **Sample/replicate** | 1 | 2 | 3 | 4 | 5 | 6 | 1 | 2 | 3 | 4 | 5 | 6 | 1 | | 2 | 3 | 4 | 5 | 6 |
| Free-living *M. conductrix* | 0.30 | 0.17 | 0.28 | 0.32 | 0.42 | 0.48 | 1.03E-09 | 7.19E-10 | 1.06E-09 | 1.09E-09 | 1.83E-09 | 1.85E-09 | 6.85E-10 | | 4.79E-10 | 7.08E-10 | 3.62E-10 | 6.10E-10 | 6.15E-10 |
| Symbiotic *M. conductrix* (1660/18) | 0.25 | 0.38 | 0.32 | 0.33 | 0.81 | 0.59 | 8.58E-09 | 1.89E-08 | 8.47E-09 | 1.16E-08 | 4.05E-08 | 1.56E-08 | 5.72E-09 | | 1.26E-08 | 5.64E-09 | 3.85E-09 | 1.35E-08 | 5.21E-09 |

|  | **cell number** | | | | | |
| --- | --- | --- | --- | --- | --- | --- |
| **Sample/replicate** | 1 | 2 | 3 | 4 | 5 | 6 |
| Free-living *M. conductrix* | 2.95E+08 | 2.95E+08 | 2.30E+08 | 2.30E+08 | 2.63E+08 | 2.63E+08 |
| Symbiotic *M. conductrix* (1660/18) | 2.88E+07 | 2.88E+07 | 2.00E+07 | 2.00E+07 | 3.75E+07 | 3.75E+07 |

| **Sample** | ** Rubisco*** pmol | ** Rubisco** pmol/cell | ** Rubisco**  (pmol/ µg prot) | ** Rubisco**  (pmol / cell) | **SD** |
| --- | --- | --- | --- | --- | --- |
| Free-living *M. conductrix* | 0.329 | 1.262E-09 | 0.151 | 5.77E-10 | 1.32E-10 |
| Symbiotic *M. conductrix* (1660/18) | 0.445 | 1.727E-08 | 0.201 | 7.75E-09 | 4.16E-09 |

**Table S6.** Rubisco quantification in free-living (*Micractinium conductrix*) and symbiotic (*M. conductrix* of *P. bursaria* CCAP1660/18) microalgae (pmol / cell) based on western blot. (See also Figure S1).

|  | *M.conductrix*  free-living  am | *M.conductrix* free-living pm | *M.conductrix* free-living D+1 am | *M.conductrix*  free-living D+1 pm | *M.conductrix* symbiont am | *M.conductrix*  symbiont pm | *M.conductrix* symbiont  D+1 am | *M.conductrix* symbiont  D+1 pm |
| --- | --- | --- | --- | --- | --- | --- | --- | --- |
|  | 0.095 | 0.206 | 0.171 | 0.256 | 0.629 | 1.328 | 0.788 | 1.660 |
|  | 0.200 | 0.454 | 0.192 | 0.303 | 0.462 | 1.389 | 0.826 | 1.544 |
|  | 0.138 | 0.288 | 0.183 | 0.226 | 0.378 | 1.498 | 0.767 | 1.150 |
| Average | 1.143 | 0.316 | 0.182 | 0.261 | 0.489 | 1.405 | 0.793 | 1.451 |
| Std.Dev. | 0.052 | 0.126 | 0.010 | 0.038 | 0.127 | 0.086 | 0.029 | 0.267 |

**Table S7**. Total starch quantification based on enzymatic assay (pg.cell^-1^) over the day (morning and afternoon) in free-living (*Micractinium conductrix*) and symbiotic (*M. conductrix* of *P. bursaria* CCAP1660/18) microalgae (three replicates).

|  | Free-living  *M. conductrix* | Symbiont  *M. conductrix* | Host  *P. bursaria* |
| --- | --- | --- | --- |
| DOC | n° of cells | n° of cells | n° of cells |
| 0 | 1.44E+05 | 193000 | 441 |
| 4 | 8.19E+06 | 558500 | 793 |
| 5 | 9.66E+06 | 1090000 | 1913 |
| 11 | 2.81E+07 | 1180000 | 1677 |

| Growth rate | | | | | |
| --- | --- | --- | --- | --- | --- |
|  | **Free-living**  ***M. conductrix*** | | **Symbiont**  ***M. conductrix*** | | **Host**  ***P. bursaria*** |
| Days |  |  | |  | |
| 0-4 | 0.042092 | 0.011068 | | 0.006117 | |
| 4-5 | 0.006878 | 0.027862 | | 0.036682 | |
| 5-11 | 0.007415 | 0.000551 | | -0.00092 | |
| 0-11 | **0.01998** | **0.00686** | | **0.00506** | |

**Table S8.** Cell counting of free-living and symbiotic *M. conductrix* and the host *Paramecium bursaria* and growth rate calculated based on the following equation: ln (final)-ln(initial)/time2-time1 (h).

| ***Chlorella vulgaris* AM time point** | Cell_1_ (*µm^3^*) | Cell_2_ (*µm^3^*) | Cell_3_ (*µm^3^*) | Cell_4_ (*µm^3^*) | Cell_5_ (*µm^3^*) | Cell_6_ (*µm^3^*) | Cell_7_ (*µm^3^*) | Cell_8_ (*µm^3^*) | **Average (*µm^3^*)** | **STD (*µm^3^*)** |
| --- | --- | --- | --- | --- | --- | --- | --- | --- | --- | --- |
| starch plates | 0.01 | 0.05 | 0.10 | 0.05 | 0.03 | 0.05 | 0.07 | 0.17 | 0.07 | 0.05 |
| starch grains | 0.00 | 0.00 | 0.08 | 0.06 | 0.00 | 0.17 | 0.00 | 0.00 | 0.04 | 0.06 |
| total starch | 0.01 | 0.05 | 0.17 | 0.11 | 0.04 | 0.22 | 0.07 | 0.17 | 0.11 | 0.08 |
| lipid droplets | 0.06 | 0.20 | 0.02 | 0.05 | 0.04 | 0.13 | 0.32 | 0.24 | 0.13 | 0.11 |
| pyrenoid | 0.05 | 0.12 | 0.10 | 0.10 | 0.07 | 0.12 | 0.12 | 0.16 | 0.11 | 0.03 |
| Cell | 5.49 | 10.26 | 7.29 | 6.75 | 5.99 | 11.48 | 14.30 | 11.16 | 9.09 | 3.16 |

| ***Chlorella vulgaris* PM time point** | Cell_1_ (*µm^3^*) | Cell_2_ (*µm^3^*) | Cell_3_ (*µm^3^*) | Cell_4_ (*µm^3^*) | Cell_5_ (*µm^3^*) | Cell_6_ (*µm^3^*) | Cell_7_ (*µm^3^*) | **Average (*µm^3^*)** | **STD (*µm^3^*)** |
| --- | --- | --- | --- | --- | --- | --- | --- | --- | --- |
| starch plates | 0.30 | 0.37 | 0.27 | 0.21 | 0.44 | 0.19 | 0.35 | 0.30 | 0.09 |
| starch grains | 0.00 | 0.02 | 0.01 | 0.04 | 0.14 | 0.01 | 0.02 | 0.03 | 0.05 |
| total starch | 0.30 | 0.39 | 0.28 | 0.26 | 0.57 | 0.19 | 0.37 | 0.34 | 0.12 |
| lipid droplets | 0.11 | 0.22 | 0.11 | 0.11 | 0.39 | 0.01 | 0.11 | 0.15 | 0.12 |
| pyrenoid | 0.16 | 0.19 | 0.17 | 0.20 | 0.10 | 0.12 | 0.18 | 0.16 | 0.04 |
| Cell | 12.63 | 15.89 | 10.88 | 11.22 | 8.14 | 9.98 | 13.46 | 11.74 | 2.52 |

| ***Micractinium conductrix***  **AM time point** | Cell_1_ (*µm^3^*) | Cell_2_ (*µm^3^*) | Cell_3_ (*µm^3^*) | Cell_4_ (*µm^3^*) | Cell_5_ (*µm^3^*) | Cell_6_ (*µm^3^*) | Cell_7_ (*µm^3^*) | Cell_8_ (*µm^3^*) | **Average (*µm^3^*)** | **STD (*µm^3^*)** |
| --- | --- | --- | --- | --- | --- | --- | --- | --- | --- | --- |
| starch plates | 0.14 | 0.14 | 0.20 | 0.11 | 0.16 | 0.08 | 0.20 | 0.30 | 0.17 | 0.07 |
| starch grains | 0.00 | 0.00 | 0.00 | 0.00 | 0.00 | 0.00 | 0.00 | 0.02 | 0.00 | 0.01 |
| total starch | 0.14 | 0.14 | 0.20 | 0.11 | 0.16 | 0.08 | 0.20 | 0.31 | 0.17 | 0.07 |
| lipid droplets | 0.04 | 0.05 | 0.07 | 0.06 | 0.05 | 0.06 | 0.05 | 0.07 | 0.05 | 0.01 |
| pyrenoid | 0.09 | 0.12 | 0.17 | 0.10 | 0.12 | 0.09 | 0.11 | 0.20 | 0.12 | 0.04 |
| Cell | 6.06 | 7.98 | 10.41 | 6.58 | 7.55 | 6.10 | 7.88 | 11.07 | 7.95 | 1.88 |

| ***Micractinium conductrix***  **PM time point** | Cell_1_ (*µm^3^*) | Cell_2_ (*µm^3^*) | Cell_3_ (*µm^3^*) | Cell_4_ (*µm^3^*) | Cell_5_ (*µm^3^*) | Cell_6_ (*µm^3^*) | Cell_7_ (*µm^3^*) | **Average (*µm^3^*)** | **STD (*µm^3^*)** |
| --- | --- | --- | --- | --- | --- | --- | --- | --- | --- |
| starch plates | 0.11 | 0.17 | 0.19 | 0.24 | 0.25 | 0.27 | 0.26 | 0.21 | 0.06 |
| starch grains | 0.00 | 0.01 | 0.00 | 0.02 | 0.02 | 0.01 | 0.00 | 0.01 | 0.01 |
| total starch | 0.11 | 0.18 | 0.19 | 0.26 | 0.27 | 0.28 | 0.27 | 0.22 | 0.06 |
| lipid droplets | 0.00 | 0.00 | 0.03 | 0.00 | 0.04 | 0.03 | 0.07 | 0.02 | 0.03 |
| pyrenoid | 0.09 | 0.11 | 0.11 | 0.16 | 0.13 | 0.18 | 0.11 | 0.13 | 0.03 |
| Cell | 5.54 | 6.57 | 6.36 | 8.22 | 7.56 | 8.08 | 9.26 | 7.37 | 1.28 |

| ***Micractinium conductrix* (isolated from *P. bursaria CCAP 1660/18)* AM** | Cell_1_ (*µm^3^*) | Cell_2_ (*µm^3^*) | Cell_3_ (*µm^3^*) | Cell_4_ (*µm^3^*) | Cell_5_ (*µm^3^*) | Cell_6_ (*µm^3^*) | Cell_7_ (*µm^3^*) | Cell_8_ (*µm^3^*) | Cell_9_ (*µm^3^*) | Cell_10_ (*µm^3^*) | **Average (*µm^3^*)** | **STD (*µm^3^*)** |
| --- | --- | --- | --- | --- | --- | --- | --- | --- | --- | --- | --- | --- |
| starch plates | 0.03 | 0.05 | 0.01 | 0.46 | 0.00 | 0.07 | 0.01 | 0.04 | 0.09 | 0.28 | 0.10 | 0.15 |
| starch grains | 0.00 | 0.00 | 0.00 | 1.16 | 0.00 | 0.00 | 0.01 | 0.00 | 0.00 | 0.00 | 0.12 | 0.37 |
| total starch | 0.03 | 0.05 | 0.01 | 1.61 | 0.00 | 0.07 | 0.01 | 0.04 | 0.09 | 0.28 | 0.22 | 0.50 |
| lipid droplets | 1.01 | 0.92 | 0.49 | 0.31 | 0.78 | 1.68 | 0.94 | 0.62 | 1.26 | 0.98 | 0.90 | 0.39 |
| pyrenoid | 1.05 | 0.92 | 0.31 | 0.11 | 0.37 | 0.56 | 0.41 | 0.34 | 1.10 | 1.27 | 0.64 | 0.40 |
| Cell | 33.67 | 29.40 | 13.34 | 15.07 | 11.88 | 21.80 | 15.67 | 14.47 | 40.19 | 39.21 | 23.47 | 11.15 |

| ***Micractinium conductrix* (isolated from *P. bursaria CCAP 1660/18)* PM** | Cell_1_ (*µm^3^*) | Cell_2_ (*µm^3^*) | Cell_3_ (*µm^3^*) | Cell_4_ (*µm^3^*) | Cell_5_ (*µm^3^*) | Cell_6_ (*µm^3^*) | Cell_7_ (*µm^3^*) | Cell_8_ (*µm^3^*) | Cell_9_ (*µm^3^*) | Cell_10_ (*µm^3^*) | Cell_11_ (*µm^3^*) | **Average (*µm^3^*)** | **STD (*µm^3^*)** |
| --- | --- | --- | --- | --- | --- | --- | --- | --- | --- | --- | --- | --- | --- |
| starch plates | 2.28 | 0.64 | 0.27 | 2.07 | 0.44 | 0.93 | 0.61 | 0.80 | 1.35 | 0.73 | 0.79 | 0.99 | 0.65 |
| starch grains | 0.95 | 0.07 | 0.03 | 1.01 | 0.04 | 0.21 | 0.10 | 0.22 | 0.52 | 0.05 | 0.05 | 0.30 | 0.37 |
| total starch | 3.22 | 0.72 | 0.30 | 3.08 | 0.47 | 1.14 | 0.72 | 1.02 | 1.87 | 0.78 | 0.84 | 1.29 | 1.01 |
| lipid droplets | 1.16 | 0.52 | 0.37 | 0.80 | 0.24 | 0.80 | 0.63 | 0.94 | 1.01 | 0.69 | 0.52 | 0.70 | 0.28 |
| pyrenoid | 1.11 | 0.42 | 0.24 | 0.98 | 0.27 | 0.67 | 0.55 | 0.61 | 1.03 | 0.71 | 0.63 | 0.66 | 0.29 |
| Cell | 34.73 | 13.76 | 8.95 | 32.55 | 9.17 | 20.48 | 17.96 | 18.41 | 30.51 | 20.45 | 17.68 | 20.42 | 8.80 |

**Table S9.** Volume (µm^3^) of carbon reserves (starch plates, starch grains and lipid droplets) based on FIB-SEM volumetrics in free-living (*Chlorella vulgaris* and *Micractinium conductrix*) and symbiotic (*M. conductrix* of *P. bursaria* CCAP1660/18) microalgae. Table shows the compartments and structures that were segmented from different datasets (in the morning=AM and in the afternoon=PM) to assess the diel carbon dynamics.

| ***Chlorella vulgaris* PM time point** | Cell_1_ (%) | Cell_2_ (%) | Cell_3_ (%) | Cell_4_ (%) | Cell_5_ (%) | Cell_6_ (%) | Cell_7_ (%) | **Average (*µm^3^*)** | **STD (*µm^3^*)** |
| --- | --- | --- | --- | --- | --- | --- | --- | --- | --- |
| starch plates | 0.023 | 0.023 | 0.025 | 0.019 | 0.054 | 0.019 | 0.026 | 0.027 | 0.012 |
| starch grains | 0.000 | 0.001 | 0.001 | 0.004 | 0.017 | 0.001 | 0.002 | 0.004 | 0.006 |
| total starch | 0.023 | 0.024 | 0.026 | 0.023 | 0.070 | 0.019 | 0.028 | 0.031 | 0.018 |
| lipid droplets | 0.009 | 0.014 | 0.010 | 0.010 | 0.048 | 0.001 | 0.008 | 0.014 | 0.015 |
| pyrenoid | 0.013 | 0.012 | 0.015 | 0.018 | 0.012 | 0.012 | 0.013 | 0.014 | 0.002 |

| ***Micractinium conductrix* AM time point** | Cell_1_ (%) | Cell_2_ (%) | Cell_3_ (%) | Cell_4_ (%) | Cell_5_ (%) | Cell_6_ (%) | Cell_7_ (%) | Cell_8_ (%) | **Average (*µm^3^*)** | **STD (*µm^3^*)** |
| --- | --- | --- | --- | --- | --- | --- | --- | --- | --- | --- |
| starch plates | 0.023 | 0.018 | 0.020 | 0.016 | 0.021 | 0.013 | 0.025 | 0.027 | 0.020 | 0.004 |
| starch grains | 0.001 | 0.000 | 0.000 | 0.000 | 0.000 | 0.000 | 0.000 | 0.002 | 0.0003 | 0.001 |
| total starch | 0.023 | 0.018 | 0.020 | 0.016 | 0.021 | 0.013 | 0.025 | 0.028 | 0.021 | 0.005 |
| lipid droplets | 0.006 | 0.006 | 0.007 | 0.009 | 0.006 | 0.009 | 0.007 | 0.006 | 0.007 | 0.001 |
| pyrenoid | 0.015 | 0.015 | 0.016 | 0.015 | 0.016 | 0.015 | 0.014 | 0.018 | 0.015 | 0.001 |

| ***Chlorella vulgaris***  **AM time point** | Cell_1_ (%) | Cell_2_ (%) | Cell_3_ (%) | Cell_4_ (%) | Cell_5_ (%) | Cell_6_ (%) | Cell_7_ (%) | Cell_8_ (%) | **Average (%)** | **STD (%)** |
| --- | --- | --- | --- | --- | --- | --- | --- | --- | --- | --- |
| starch plates | 0.002 | 0.005 | 0.014 | 0.007 | 0.006 | 0.004 | 0.005 | 0.015 | 0.007 | 0.005 |
| starch grains | 0.000 | 0.000 | 0.010 | 0.009 | 0.000 | 0.015 | 0.000 | 0.000 | 0.004 | 0.006 |
| total starch | 0.002 | 0.005 | 0.024 | 0.016 | 0.006 | 0.019 | 0.005 | 0.015 | 0.011 | 0.008 |
| lipid droplets | 0.011 | 0.020 | 0.003 | 0.007 | 0.006 | 0.011 | 0.022 | 0.022 | 0.013 | 0.007 |
| pyrenoid | 0.009 | 0.012 | 0.014 | 0.015 | 0.012 | 0.010 | 0.008 | 0.014 | 0.012 | 0.003 |

| ***Micractinium conductrix***  **PM time point** | Cell_1_ (%) | Cell_2_ (%) | Cell_3_ (%) | Cell_4_ (%) | Cell_5_ (%) | Cell_6_ (%) | Cell_7_ (%) | **Average (*µm^3^*)** | **STD (*µm^3^*)** |
| --- | --- | --- | --- | --- | --- | --- | --- | --- | --- |
| starch plates | 0.020 | 0.025 | 0.029 | 0.029 | 0.033 | 0.033 | 0.028 | 0.028 | 0.004 |
| starch grains | 0.000 | 0.002 | 0.000 | 0.002 | 0.002 | 0.002 | 0.000 | 0.001 | 0.001 |
| total starch | 0.021 | 0.027 | 0.029 | 0.032 | 0.036 | 0.034 | 0.029 | 0.030 | 0.005 |
| lipid droplets | 0.000 | 0.000 | 0.005 | 0.000 | 0.006 | 0.003 | 0.008 | 0.003 | 0.003 |
| pyrenoid | 0.016 | 0.016 | 0.018 | 0.020 | 0.018 | 0.022 | 0.012 | 0.017 | 0.003 |

| ***Micractinium conductrix* (isolated from *P. bursaria CCAP 1660/18)* AM** | Cell_1_ (%) | Cell_2_ (%) | Cell_3_ (%) | Cell_4_ (%) | Cell_5_ (%) | Cell_6_ (%) | Cell_7_ (%) | Cell_8_ (%) | Cell_9_ (%) | Cell_10_ (%) | **Average (%)** | **STD (%)** |
| --- | --- | --- | --- | --- | --- | --- | --- | --- | --- | --- | --- | --- |
| starch plates | 0.001 | 0.002 | 0.000 | 0.030 | 0.000 | 0.003 | 0.000 | 0.003 | 0.002 | 0.007 | 0.005 | 0.009 |
| starch grains | 0.000 | 0.000 | 0.000 | 0.077 | 0.000 | 0.000 | 0.001 | 0.000 | 0.000 | 0.000 | 0.008 | 0.024 |
| total starch | 0.001 | 0.002 | 0.000 | 0.107 | 0.000 | 0.003 | 0.001 | 0.003 | 0.002 | 0.007 | 0.013 | 0.033 |
| lipid droplets | 0.030 | 0.031 | 0.037 | 0.021 | 0.066 | 0.077 | 0.060 | 0.043 | 0.031 | 0.025 | 0.042 | 0.019 |
| pyrenoid | 0.031 | 0.031 | 0.023 | 0.007 | 0.031 | 0.025 | 0.026 | 0.024 | 0.027 | 0.032 | 0.026 | 0.007 |

| ***Micractinium conductrix* (isolated from *P. bursaria CCAP 1660/18)* PM** | Cell_1_ (%) | Cell_2_ (%) | Cell_3_ (%) | Cell_4_ (%) | Cell_5_ (%) | Cell_6_ (%) | Cell_7_ (%) | Cell_8_ (%) | Cell_9_ (%) | Cell_10_ (%) | Cell_11_ (%) | **Average (%)** | **STD (%)** |
| --- | --- | --- | --- | --- | --- | --- | --- | --- | --- | --- | --- | --- | --- |
| starch plates | 0.066 | 0.047 | 0.030 | 0.063 | 0.047 | 0.045 | 0.034 | 0.043 | 0.044 | 0.036 | 0.045 | 0.045 | 0.011 |
| starch grains | 0.027 | 0.005 | 0.003 | 0.031 | 0.004 | 0.010 | 0.006 | 0.012 | 0.017 | 0.002 | 0.003 | 0.011 | 0.010 |
| total starch | 0.093 | 0.052 | 0.033 | 0.095 | 0.052 | 0.056 | 0.040 | 0.056 | 0.061 | 0.038 | 0.048 | 0.057 | 0.020 |
| lipid droplets | 0.033 | 0.038 | 0.041 | 0.025 | 0.026 | 0.039 | 0.035 | 0.051 | 0.033 | 0.034 | 0.030 | 0.035 | 0.007 |
| pyrenoid | 0.032 | 0.031 | 0.027 | 0.030 | 0.029 | 0.033 | 0.031 | 0.033 | 0.034 | 0.035 | 0.035 | 0.032 | 0.003 |
| Cell |  |  |  |  |  |  |  |  |  |  |  |  |  |

**Table S10.** Volume occupancy (%) of carbon storage (starch and lipid droplets) in free-living (*Chlorella vulgaris* and *Micractinium conductrix*) and symbiotic (*M. conductrix* of *P. bursaria* CCAP1660/18) microalgae. Table shows the volume occupancy (%) that each structure/compartment occupies in the cell in the morning (AM) and in the afternoon (PM).

|  | ***M.conductrix* free-living am** | ***M.conductrix* free-living pm** | ***M.conductrix* free-living D+1 am** | ***M.conductrix* free-living D+1 pm** | ***M.conductrix* symbiont am** | ***M.conductrix* symbiont pm** | ***M.conductrix* symbiont D+1 am** | ***M.conductrix* symbiont *D+1 pm*** | ***M.conductrix* symbiont D+3 am** | ***M.conductrix* symbiont D+3 pm** |
| --- | --- | --- | --- | --- | --- | --- | --- | --- | --- | --- |
|  | 0.018 | 0.012 | 0.017 | 0.022 | 0.021 | 0.015 | 0.023 | 0.035 | 0.039 | 0.041 |
|  | 0.016 | 0.012 | 0.020 | 0.014 | 0.014 | 0.018 | 0.034 | 0.025 | 0.032 | 0.036 |
|  | 0.019 | 0.013 | 0.018 | 0.018 | 0.023 | 0.016 | 0.030 | 0.036 | 0.035 | 0.036 |
|  | 0.017 | 0.016 | 0.021 | 0.016 | 0.019 | 0.016 | 0.034 | 0.036 | 0.037 | 0.034 |
|  | 0.016 | 0.015 | 0.021 | 0.014 | 0.014 | 0.021 | 0.026 | 0.037 | 0.032 | 0.039 |
|  | 0.035 | 0.024 | 0.018 | 0.021 | 0.019 | 0.017 | 0.022 | 0.053 | 0.034 | 0.035 |
|  | 0.031 | 0.025 | 0.016 | 0.016 | 0.021 | 0.020 | 0.025 | 0.039 | 0.043 | 0.034 |
|  | 0.033 | 0.039 | 0.019 | 0.019 | 0.018 | 0.024 | 0.034 | 0.043 | 0.035 | 0.034 |
|  | 0.027 | 0.022 | 0.018 | 0.017 | 0.022 | 0.022 | 0.020 | 0.042 | 0.034 | 0.037 |
|  | 0.031 | 0.024 | 0.031 | 0.013 | 0.025 | 0.018 | 0.030 | 0.043 | 0.044 | 0.039 |
|  | 0.017 | 0.012 | 0.017 | 0.016 | 0.016 | 0.022 | 0.025 | 0.026 | 0.050 | 0.038 |
|  | 0.016 | 0.012 | 0.017 | 0.014 | 0.012 | 0.021 | 0.035 | 0.025 | 0.048 | 0.026 |
|  | 0.017 | 0.011 | 0.016 | 0.015 | 0.014 | 0.019 | 0.031 | 0.026 | 0.062 | 0.046 |
|  | 0.015 | 0.012 | 0.017 | 0.017 | 0.015 | 0.024 | 0.024 | 0.026 | 0.057 | 0.044 |
|  | 0.018 | 0.014 | 0.015 | 0.016 | 0.012 | 0.015 | 0.023 | 0.024 | 0.052 | 0.039 |
| **Average** | 0.022 | 0.017 | 0.019 | 0.016 | 0.017 | 0.019 | 0.028 | 0.034 | 0.042 | 0.037 |
| **Std.Dev.** | 0.007 | 0.007 | 0.004 | 0.003 | 0.004 | 0.003 | 0.005 | 0.009 | 0.010 | 0.005 |

**Table S11.** Lipid droplets quantification based on fluorescence of Nile red in free-living (*Chlorella vulgaris* and *Micractinium conductrix*) and symbiotic (*M. conductrix* of *P. bursaria* CCAP1660/18) microalgae (arbitrary units per cell).

| **Sample** | **Isotope** | **condition / time** | **Replicate** | **Number of cells** | **d 13C/12C (‰)** | **µg C** | ** d13C/12C(‰)** | ** %C** |
| --- | --- | --- | --- | --- | --- | --- | --- | --- |
| *Paramecium bursaria* (CCAP1660/18) | n/n | control | 1 | 9150 | -27.77 | 62.75 | -27.64 | 59.41 |
|  | n/n | control | 2 | 9150 | -27.52 | 56.08 |  |  |
|  | 13C-glucose | light_24h | 1 | 6975 | 9624.82 | 35.92 | 9565.51 | 35.75 |
|  | 13C-glucose | light_24h | 2 | 6975 | 9506.20 | 35.57 |  |  |

**Table S12**. ^13^C-glucose uptake by the host *Paramecium bursaria* (CCAP1660/18) in symbiosis with *M. conductrix*

| **Total starch (pg.cell^-1^)** | | | | |
| --- | --- | --- | --- | --- |
| **cell** | **Control T0h** | **Control T6h** | **Glucose T0h** | **Glucose T6h** |
|  | 0.574 | 5.113 | 0.982 | 2.126 |
| *P. bursaria* (CCAP1660/18) | 1.322 | 3.859 | 1.521 | 2.832 |
|  | 0.364 | 3.850 | 0.433 | 2.603 |
| **Average** | 0.753 | 4.274 | 0.978 | 2.520 |
| **SD** | 0.504 | 0.727 | 0.544 | 0.360 |

**Table S13.** Total starch quantification of *Paramecium bursaria* in symbiosis with the microalga *Micractinium conductrix* when exposed to 75 mM of glucose and the control condition (no glucose). Host cells were harvested at T0h=9.30 am and T6h=3.30 pm. Starch quantification is expressed as pg.cell^-1^.
